# Functional Architecture of Executive Control and Associated Event-Related Potentials

**DOI:** 10.1101/2021.01.30.428901

**Authors:** Amirsaman Sajad, Steven P. Errington, Jeffrey D. Schall

## Abstract

Medial frontal cortex enables executive control by monitoring relevant information and using it to adapt behavior. In macaques performing a saccade countermanding (stop-signal) task, we recorded EEG over and neural spiking across all layers of the supplementary eye field (SEF). We report the laminar organization of concurrently activated neurons monitoring the conflict between incompatible responses and the timing of events serving goal maintenance and executive control. We also show their relation to coincident event-related potentials (ERP). Neurons signaling response conflict were largely broad-spiking found across all layers. Neurons signaling the interval until specific task events were largely broad-spiking neurons concentrated in L3 and L5. Neurons predicting the duration of control and sustaining the task goal until the release of operant control were a mix of narrow- and broad-spiking neurons confined to L2/3. We complement these results with the first report of a monkey homologue of the N2/P3 ERP complex associated with response inhibition. N2 polarization varied with error likelihood and P3 polarization varied with the duration of expected control. The amplitude of the N2 and P3 were predicted by the spike rate of different classes of neurons located in L2/3 but not L5/6. These findings reveal important, new features of the cortical microcircuitry supporting executive control and producing associated ERP.

---

Effective control of behavior is necessary to achieve goals, especially when faced with competing instructions inducing response conflict and requiring inhibition of prepotent responses and maintenance of task goals, and adaptation of performance. These features of executive control are investigated with the countermanding (stop-signal) task ^1^, during which macaque monkeys, like humans, exert response inhibition and adapt performance based on stimulus history, response outcomes, and the temporal structure of task events ^2^.

Medial frontal cortex enables executive control, but circuit-level mechanisms remain uncertain ^3, 4^. Hypotheses on executive control function have been tested in humans using noninvasive ERP measures derived from a negative-positive waveform known as the N2/P3 associated with stopping ^5^. However, their cortical source is unknown. Mechanistic hypotheses about the basis of these signals require information about neural spiking patterns across cortical layers ^6^. Moreover, understanding function at the resolution of layers can clarify circuit-level mechanisms because neurons in different layers have different extrinsic anatomical connections. We can obtain such information from the supplementary eye field, an agranular area on the dorsomedial convexity in macaques, immediately beneath where the frontal ERPs are sampled. SEF contributes to proactive but not reactive inhibition ^7^ and its activation improves performance in the countermanding task by delaying response time^8^ through postponing the accumulation of pre-saccadic activity ^9^. SEF also supports working memory ^10, 11^, and signals surprise ^12^, event timing ^13, 14^, response conflict ^15^, plus errors and reinforcement ^16^. SEF in macaques is homologous to SEF in humans ^17^.

The canonical cortical microcircuit derived from granular sensory areas ^18^ does not explain agranular frontal areas like SEF ^19, 20, 21, 22, 23^. Recently we described the laminar microcircuitry of performance monitoring signals in the SEF, and relationship to the ERP indexing error monitoring known as the error-related negativity (ERN)^16^. Here we describe the laminar microcircuitry of signals that monitor events occurring during successful stopping performance. We define three classes of neurons that concurrently signal response conflict, timing of events, and maintenance of task goals. We also establish that macaque monkeys produce the N2/P3 ERP associated with response inhibition, elucidating task factors indexed by this ERP complex and the neuron classes predicting their polarization.

## RESULTS

### Countermanding performance, neural sampling, and functional classification

Neurophysiological and electrophysiological data was recorded from two macaque monkeys performing the saccade countermanding task with explicit feedback tone cues (**Fig. 1a**) ^24^. Data collection and analysis was informed by the consensus guide for the stop-signal task ^25^. In 29 sessions we acquired 33,816 trials (Monkey Eu, male, 12 sessions 11,583 trials; X, female, 17 sessions 22,233 trials). Typical performance was produced by both monkeys. Response times (RT) on failed inhibition trials (noncanceled trials) (mean ± SD Eu: 294 ± 179 ms; X: 230 ± 83 ms) were systematically shorter than those on no stop-signal trials (Eu: 313 ± 119 ms, X: 263 ± 112 ms; mixed effects linear regression grouped by monkey, t(27507) = −17.4, p < 10^-5^) (**Fig. 1b**). Characteristically, the probability of noncanceled errors increased with stop-signal delay (SSD) (**Fig. 1b**). These two observations validate the use of the independent race model ^26^ to estimate the stop-signal reaction time (SSRT), the time needed to cancel a partially prepared saccade. Accordingly, neural modulation before SSRT can contribute to stopping but that after SSRT cannot ^7, 26^. SSRT across sessions (Eu: 118 ± 23 ms, X: 103 ± 24 ms) did not differ between monkeys (t(27) = −1.69, p = 0.1025). While there were other classes of errors made in the task, they were infrequent and therefore inconsequential to this study. Therefore, P(error) refers to the probability of noncanceled errors.

**Fig. 1.**
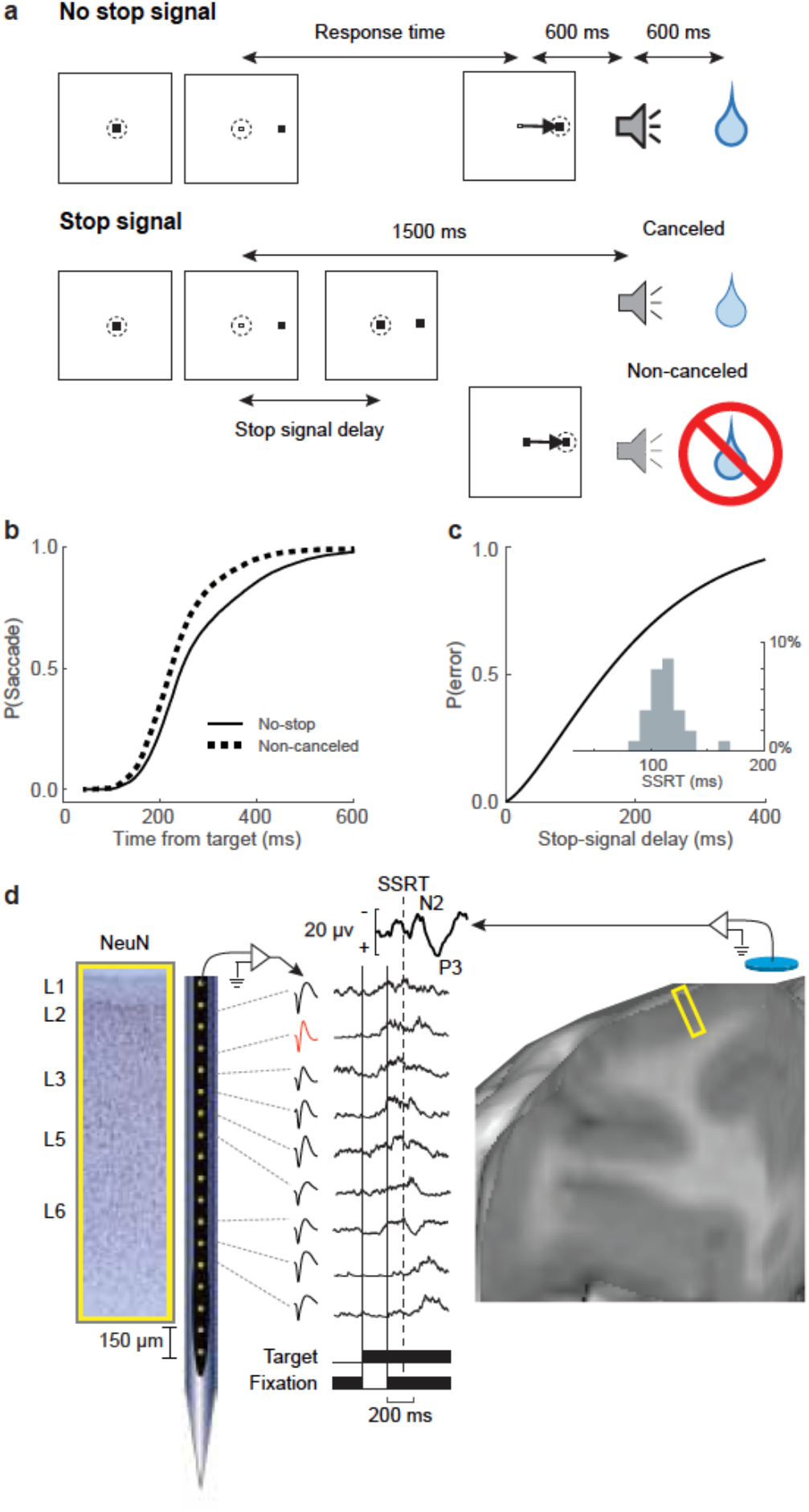
Experimental approach. **a**, Saccade countermanding task. Monkeys initiated trials by fixating a central point. After a variable time, the center of the fixation point was extinguished, and a peripheral target was presented at one of two possible locations. On no stop-signal trials monkeys were required to shift gaze to the target, whereupon after 600 ± 0 ms a high-pitch auditory feedback tone was delivered, and 600 ± 0 ms later fluid reward was provided. On stop-signal trials (∼40% of trials) after the target appeared, the center of the fixation point was re-illuminated after a variable stop-signal delay, which instructed the monkey to cancel the saccade in which case the same high-pitch tone was presented 1,500 ± 0 ms after target presentation followed 600 ± 0 ms later by fluid reward. Stop-signal delay was adjusted such that monkeys successfully canceled the saccade in ∼50% of trials. In the remaining trials, monkeys made non-canceled errors, which were followed after 600 ± 0 ms by a low-pitch tone, and no reward was delivered. Monkeys could not initiate trials earlier after errors. **b**, Grand average cumulative distributions of all RT for both monkeys on trials with no stop-signal (solid) and non-canceled errors (dashed). **c**, Grand average probability of non-canceled errors (P(error)) as a function of stop-signal delay. Inset shows the distribution of SSRT across all sessions for both monkeys. **d**, Neural spiking was recorded across all layers of agranular SEF (NeuN stain) using Plexon U-probe. Neurons with both broad (black) and narrow (red) spikes were sampled. Spiking modulation was measured relative to presentation of task events (thin solid, visual target; thick solid, stop-signal) and performance measures like SSRT (dashed vertical). Simultaneously, EEG was recorded from the cranial surface with an electrode positioned over the medial frontal cortex (10-20 location Fz). Yellow rectangle portrays cortical area sampled in a T1 MR image.

EEG was recorded with leads placed on the cranial surface beside the chamber over medial frontal cortex while a linear electrode array (Plexon, 24 channels, 150 µm spacing) was inserted in SEF (**Fig. 1c**). SEF was localized by anatomical landmarks and intracortical electrical microstimulation ^20^. We recorded neural spiking in 29 sessions (Eu: 12, X: 17) sampling activity from 5 neighboring sites. Overall, 575 single units (Eu: 244, X: 331) were isolated, of which 213 (Eu: 105, X: 108) were modulated after SSRT. The description of the laminar distribution of signals is based on 16 of the 29 sessions during which electrode arrays were oriented perpendicular to cortical layers and we could assign neurons to different layers confidently ^20^ (see Supplementary Fig. 1 of ^16^). Additional information about laminar structure was assessed through the pattern of phase-amplitude coupling across SEF layers ^22^. Due to variability in the estimates and the indistinct nature of the L6 border with white matter, some units appeared beyond the average gray-matter estimate; these were assigned to the nearest cellular layer. In all, 119 isolated neurons (Eu: 54; X: 65) contributed to the results on laminar distribution of executive control signals subserving successful stopping (**Supplementary Table 1a)**.

To identify neural activity associated with saccade countermanding, we examined the activity across different SSDs on canceled trials in which the subject successfully inhibited the movement, and latency-matched no stop-signal trials in which no stopping was required ^27^. A consensus cluster algorithm ^28^ with manual curation identified neurons with response facilitation (n = 129) and response suppression (n = 84) following the stop-signal (**Supplementary Figure 1**). Simultaneously, we observed distinct patterns in the cranial EEG related to successful stopping with characteristic N2 and P3 components (**Fig 1c**). Whilst we previously described neural signals after errors and associated with reward, here we focused on the interval in which response inhibition was accomplished. Specifically, we quantified spiking before and after SSRT and before the feedback tone (T_tone_), which terminated operant control on behavior. To elucidate contributions of the diverse neurons, we compared and contrasted how well neural spiking related to a variety of computational parameters inherent in the task.

First, performance of the stop-signal task is explained as the outcome of a race between stochastic GO and STOP processes ^26^, instantiated by specific interactions enabling the interruption of the GO process by a STOP process ^29, 30^ (**Supplementary Figure 2a**). An influential theory of medial frontal function posits that it encodes the conflict between mutually incompatible processes ^31^. Such conflict arises naturally as the mathematical product of the activation of GO and STOP units, which is proportional to P(error). Hence, neural signals that scale with P(error) can encode conflict in this task.

Second, inspired by reinforcement learning models, we considered the possibility that neural signals reflect the error-likelihood associated with an experienced SSD ^32^. Note, on some stop-signal error trials, the response was generated before the stop-signal appeared. The error-likelihood can only form based on trials in which SSD elapsed before RT such that monkeys could see the stop-signal (referred to as SS_seen_). Hence, neural signals that scale with P(error | SS_seen_) can encode error likelihood in this task. Conflict indexed by P(error) and error likelihood indexed by P(error | SS_seen_) diverge at longer SSDs (**Supplementary Figure 2c**).

Third, monkeys can learn the timing of the various task events (**Supplementary Figure 2b**). For example, monkeys are sensitive to the adjustments of SSD that are made to maintain ∼50% success on stop-signal trials ^33^. Previous research has characterized time perception ^34, 35, 36^. Key features include sensitivity to log(interval) versus its absolute value with precision decreasing with duration and sensitivity to instantaneous expectation (i.e., hazard rate) of events (**Supplementary Figure 2d-e**). Therefore, neural activity around the time of SSD can scale with the timing or expectation of the stop-signal ^13, 14, 37, 38^. This expectation can be derived from experienced SSD and the estimated probability of stop-signal appearance (**Supplementary Figure 2e**). Moreover, to earn reward, monkeys were required to maintain fixation on the target on trials with no stop-signal or on the fixation spot on canceled trials for an extended period (T_tone_) until a tone secondary reinforcer (feedback) announced delivery of reward after another interval. Hence, neural activity associated with the tone can scale with the timing or instead the expectation of the tone, which was variable on canceled trials but predictable based on the experienced SSD (**Supplementary Figure 2d**).

Alternatives were compared through mixed-effects model-comparison with Bayesian Information Criteria (BIC). As detailed below, many neurons signaled conflict and more signaled event timing with activity sustained until earning reward.

### Monitoring Conflict

We found 75 neurons in SEF with transient facilitation after SSRT on canceled trials, compared to latency-matched no stop-signal trials, that was proportional to P(error) **(Fig. 2; Supplementary Figure 3; Supplementary Table 2)**. The transient modulation in these neurons was not just a visual response to the stop-signal because it did not happen on noncanceled trials (**Supplementary Figure 1e**). On average, this modulation started 99 ± 8 ms (mean ± SEM) after SSRT. **Figure 2a** shows the recruitment of these neurons through time. Nearly all (71/75) were recruited after SSRT, and the proportion of recruited neurons peaked at ∼60% ∼110 ms after SSRT and gradually reduced to 8% after 500 ms (**Fig. 2a**). As this facilitation occurs after SSRT, it cannot contribute to reactive response inhibition ^7^. On canceled trials a minority of these neurons produced weak, persistent activity that lasted until the tone, and some also exhibited a brief transient response following the tone (**Fig. 2; Supplementary Figure 1c**).

**Fig. 2.**
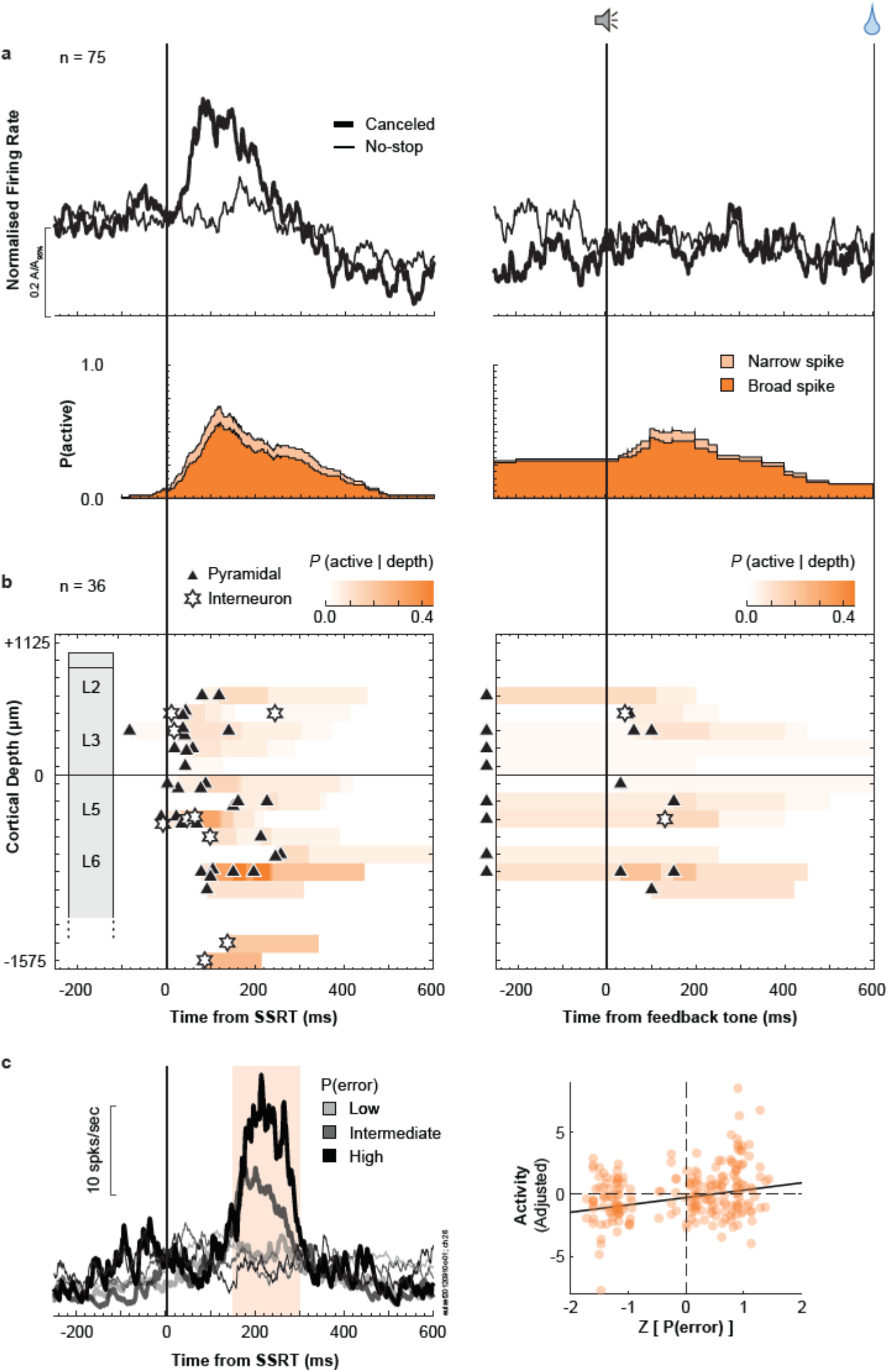
Time-depth organization of Conflict neuron spiking in SEF. **a,** Normalized population response of neurons with transient facilitation in discharge rate on successfully canceled (thick) relative to latency-matched no stop-signal (thin) trials for early SSD (top). Recruitment of this signal through time relative to SSRT (left) and auditory feedback tone (right), with dark and light shades representing the recruitment of broad-spiking (spike width ≥ 250 µs) and narrow-spiking (< 250 µs) neurons (bottom). Recruitment on SSRT-aligned activity (left panel) is defined as the difference between canceled and no stop-signal trials. Recruitment on tone-aligned activity (right panel) is defined as the activity on canceled trials relative to the baseline. Modulations starting 300ms after the tone are not included. **b,** Time-depth plot showing latency and proportion of recruited neurons through time at each depth from perpendicular penetrations. Symbols mark beginning of modulation for broad-spiking neurons (black triangles) and narrow-spiking neurons (white stars). Color map indicates the percentage of neurons relative to the overall sampling density (**Supplementary Figure 1a**) producing this signal through time at each depth. Dashed horizontal line marks L3-L5 boundary. The lower boundary of L6 is not discrete. **c (left),** Comparison of response of a representative neuron on successfully canceled (thick) relative to latency-matched no stop-signal (thin) trials for low (lighter) and higher (darker) P(error). Shaded area represents significant difference in discharge rate between the two conditions. **c (right)** Relationship between spike rate, sampled from the period with significant modulation for each neuron and the corresponding P(error). Along the ordinate scale is plotted the spiking rate, adjusted for neuron-specific variations. Along the abscissa scale is plotted the normalized P(error) (z-scale). In all, 225 points are plotted. Variation of spiking rate was best predicted by P(error) (highlighted by the best-fit line; **Supplementary Table 2**).

We assessed how the magnitude of this transient modulation after SSRT varied with the various task and performance parameters described above. The magnitude of this modulation varied most closely with P(error) – a measure of conflict (Mixed-effects linear regression grouped by neuron, t(104) = 3.57, p = 5.4×10^-4^). This conflict model obtained lower BIC than models of SSD or any other quantity, with weak support against the P(error | SS_seen_) (ΔBIC = 1.29) and strong support against other models (ΔBIC > 2.7) (**Fig 2c; Supplementary Table 2**). In contrast, the spike rate immediately before the feedback tone was unrelated to any factor related to its time or anticipation. Henceforth, we refer to these as Conflict neurons.

We noted that the vast majority (65/75) of Conflict neurons did not signal noncanceled errors, supporting previous findings (**Supplementary Table 3**) ^15, 16^. However, many (41/75) also exhibited modulation that signaled outcome following the feedback tone and around the time of reward. Some exhibited higher discharge rates on unrewarded trials (previously identified as Loss signal ^16^), and some, higher discharge rates on rewarded trials (previously identified as Gain signal ^16^). The multiplexing of the conflict monitoring signal with Gain and Loss signals (in different task epochs) did not differ from that predicted based on their sampling prevalence (*X*^2^ (3, *N* = 575) = 1.02, p = 0.79; **Supplementary Table 3**).

Conflict neurons were found at all recording sites but more commonly at some (*X*^2^ (4, *N* = 575) = 11.6, p = 0.020). Using trough-to-peak duration of the action potential waveform, the majority (63/75) had broad spikes consistent with pyramidal neurons. This distribution did not differ from the overall sampling distribution in SEF (*X*^2^ (1, *N* = 575) = 0.67, p = 0.41).

From sessions with perpendicular penetrations, we assigned 36 of the 75 Conflict neurons to a cortical layer. They were found in all layers at a relative prevalence across layers indistinguishable from that of the overall sampling distribution (*X*^2^ (4, *N* = 293) = 4.28, *p* = 0.37; **Fig 2b; Supplementary Table 1b**). The timing of the modulation did not differ between L2/3 and L5/6 (t(34) = 0.3367, p = 0.74, two tailed). The few neurons modulating with the tone were observed sparsely across all layers.

In summary, as reported previously ^15^, neurons in SEF modulate in a manner consistent with signaling the co-activation of gaze-shifting (GO) and gaze-holding (STOP) processes. This co-activation has previously been interpreted as conflict ^31, 39^ The new results show that these neurons are distributed across all SEF layers and are predominantly putative pyramidal neurons with broad spikes.

### Time keeping

Monkeys adapt performance by learning the temporal regularities of the task ^33, 40^. We identified neurons across the layers of SEF with modulation representing event timing and interval duration through facilitation, suppression, and ramping activity ^13, 14, 37, 41^ (**Supplementary Fig 2c, d**). Following target presentation, the discharge rate of many neurons (N = 84) ramped up until the saccade on trials in which they were generated (no stop-signal or noncanceled error trials). On canceled trials, however, the discharge rate was instead abruptly reduced after SSRT (**Fig 3a; Supplementary Fig 1c-e**). Because the first pronounced suppression began after SSRT, these neurons cannot contribute directly to response inhibition. Relative to SSRT, these neurons were suppressed before the facilitation in the conflict monitoring neurons (t-test, t(157) = −3.60, p = 4.2×10^-4^). The ramping activity from target to SSRT varied best with the time-based models of SSD (t(250) = 12.62, p = 0.0013) with strong support against other models (Δ BIC > 2.7) (**Supplementary Table 2**). The log-transformed model outperformed the linear model but evidence against the linear model was weak (ΔBIC = 1.35). Because the discharge rate dropped sharply on canceled trials but not on noncanceled stop-signal trials (**Supplementary Fig 1e**), we conjecture that these neurons encode the temporal aspects of events leading to successful stopping and not the timing of the stop-signal appearance per se. Once successful stopping occurred, these neurons were suppressed.

**Fig. 3.**
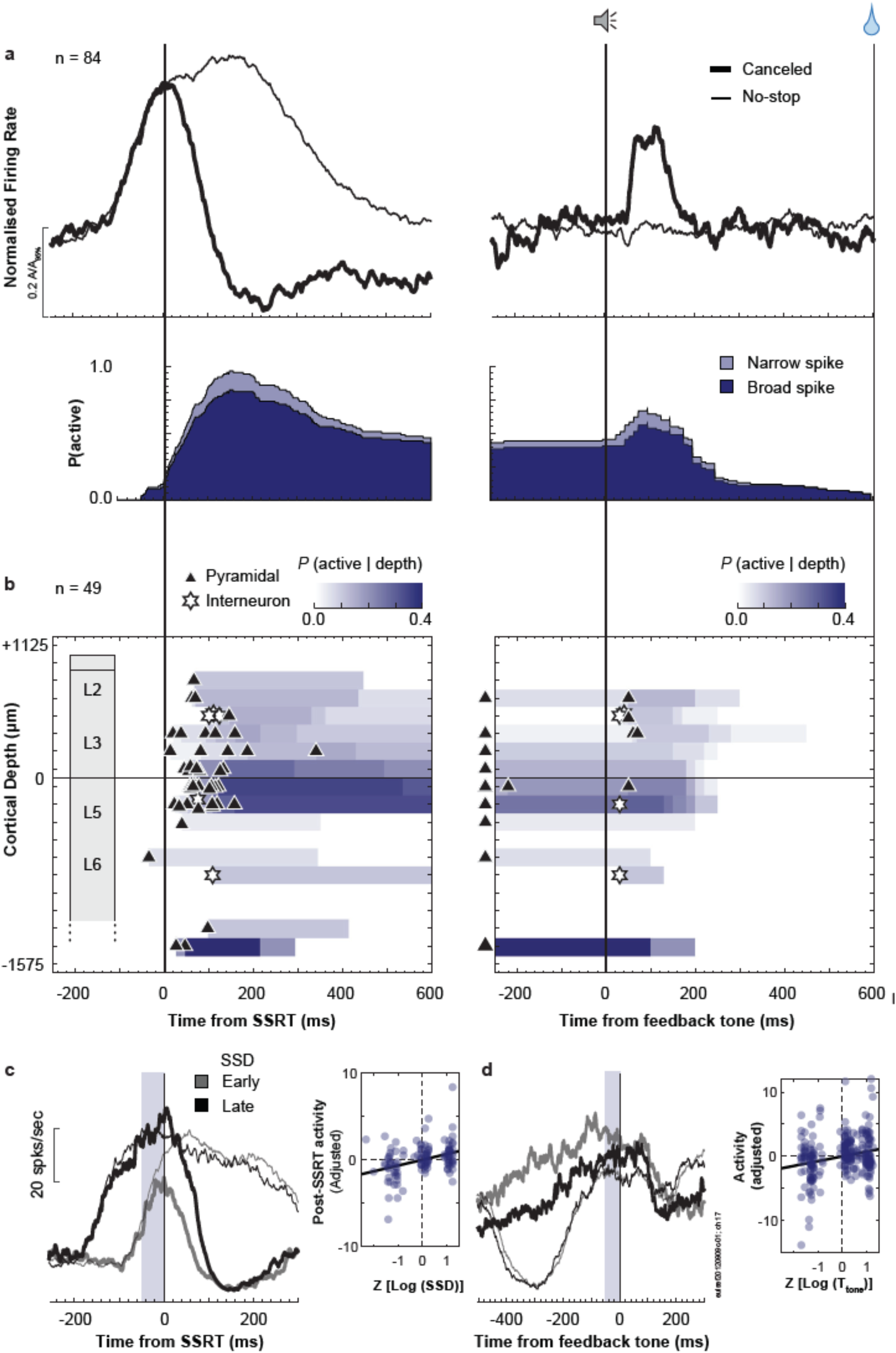
Time-depth organization of Event Timing neuron spiking in SEF. **a,** Normalized population response of neurons with suppression of discharge rate on successfully canceled (thick) relative to latency-matched no stop-signal (thin) trials for early SSD (bottom). Recruitment of signal through time relative to SSRT (left) and auditory feedback tone (right), with dark and light shades representing the recruitment of broad-spiking (spike width ≥ 250 µs) and narrow-spiking (< 250 µs) neurons (bottom). Recruitment on SSRT-aligned activity (left panel) is defined as the difference between canceled and no stop-signal trials. Recruitment on tone-aligned activity (right panel) is defined as the activity on canceled trials relative to the baseline. Modulations starting 300ms after the tone are not shown. **b,** Time-depth plot showing latency and proportion of recruited neurons through time at each depth from perpendicular penetrations. Symbols mark beginning of modulation for broad-spiking neurons (black triangles) and narrow-spiking neurons (white stars). Color map indicates the percentage of neurons relative to the overall sampling density (**Supplementary Figure 1a**) producing this signal through time at each depth. Dashed horizontal line marks L3-L5 boundary. The lower boundary of L6 is not discrete. **c,** Left panel shows response of a representative neurons on successfully canceled (thick) and latency-matched no stop-signal (thin) trials for early (lighter) and later (darker) SSD. Pre-SSRT ramping activity occurs irrespective of trial class. Shaded area represents the time epoch used for sampling neuron activity (50 ms window pre-SSRT). Right panel plots **r**elationship between discharge rate in the sampling interval and stop-signal delay. Along the ordinate scale is plotted the normalized spiking rate, adjusting for neuron-specific variations. Along the abscissa scale is plotted the normalized (z-transformed) stop-signal delay in logarithmic scale. In all, 252 points (84 neurons) are plotted. Each point plots the average spike-density and associated Log (SSD) in one of 3 bins corresponding to early-, mid-, or late-SSD, for each neuron. Variation of spiking rate was best predicted by the time of the stop-signal (highlighted by best-fit line). **d,** Left panel plots response of the same representative neuron as **c** indicating pre-tone ramping activity on successfully canceled (thick) relative to latency-matched no stop-signal (thin) trials for early (lighter) and later (darker) SSD. Shaded area represents the time epoch used for sampling neuron activity (50 ms window pre-Tone). Right panel plots relationship between discharge rate in the sampling interval and the time of feedback relative to stop-signal. Along the ordinate scale is plotted the spiking rate, adjusted for neuron-specific variations. Along the abscissa scale is plotted the normalized stop-signal delay in logarithmic scale (z-scale). In all, 144 points (38 neurons with pre-tone activity on canceled trials) are plotted. Each point plots the average spike-density and associated log (feedback time) in one of 3 bins corresponding to early-, mid-, or late-SSD, for each neuron. Variation of spiking rate was best predicted by the time of the feedback time (highlighted by best-fit line; **Supplementary Table 2**).

A subset of these neurons (29/84) also exhibited monotonic ramping of discharge rate following the sharp suppression, persisting until after the feedback tone whereupon the spike rate again decreased (**Fig 3d**). In some neurons this decrease followed a brief transient response (**Fig 3a**). The variation in dynamics of the ramping before the tone was best accounted for by the time of the feedback tone after the stop-signal (t(112) = 3.41, 9.1×10^-4^) with strong support against other models (Δ BIC > 5.0). The linear and log-transformed models were indistinguishable (ΔBIC < 0.1) (**Supplementary Table 2**). The termination of this modulation was best described by the time of the feedback tone and not the time at which fixation from stop-signal was broken (**Supplementary Figure 4c).**

Because the ramping activity in this population of neurons scaled with the time of the stop-signal and the tone, followed by immediate suppression after their occurrence, we conjecture that these neurons represent event timing to accomplish the task. We will refer to these neurons as Event Timing neurons. While all of these neurons encoded the timing of events related to successful stopping, only ∼30% also encoded the timing of the feedback tone.

Event Timing neurons were found in all penetrations, but more commonly in certain sites (*X*^2^ (4, N = 575) > 39.3, p < 10^-5^) (**Fig 3b, Supplementary Table 1a**). The majority (73 / 84) had broad spikes, corresponding to the overall sampling distribution in SEF (*X*^2^ (1, *N = 575*) = 2.56, p = 0.11). From sessions with perpendicular penetrations, we assigned the layer of 49 of the 84 neurons. The laminar organization of these neurons did not differ from the overall laminar sampling distribution (*X*^2^ (4, *N* = 293) = 7.33, p = 0.12). However, those with ramping activity before the tone (which resulted in a prolonged differential activity level between no-stop and canceled trials) were more confined to lower L3 and upper L5. The time of modulation after SSRT or around the tone did not vary across layers.

In summary, neurons in SEF exhibit ramping activity that can signal the time preceding critical events for successful task performance. The new results show that these neurons are distributed across all SEF layers and are predominantly pyramidal neurons. Often these neurons also exhibited post-feedback ramping activity leading to the time of reward delivery. Accordingly, a higher proportion of these neurons were identified as Gain neurons compared to that predicted by the prevalence of Gain and Loss neurons^16^ (*X*^2^ (3, *N* = 575) = 44.86, p = < 10^-5^; **Supplementary Table 3**).

### Goal Maintenance

By design, to earn reward on canceled trials, monkeys were required to maintain fixation on the stop-signal until an auditory feedback tone occurred. As such, the state of response inhibition needed to be maintained for an arbitrary interval. Many other neurons (N = 54) in SEF produced spike rate modulation sufficient to contribute to this maintenance (**Fig 4**). These neurons produced significantly greater discharge rates on canceled trials after SSRT, compared to latency-matched no-stop trials. Modulation was weak or absent on noncancelled error trials, so this activity was not a response to the stop-signal. This modulation began too late to contribute to response inhibition but persisted while fixation maintenance was required (**Supplementary Figure 1d, e**).

**Fig. 4.**
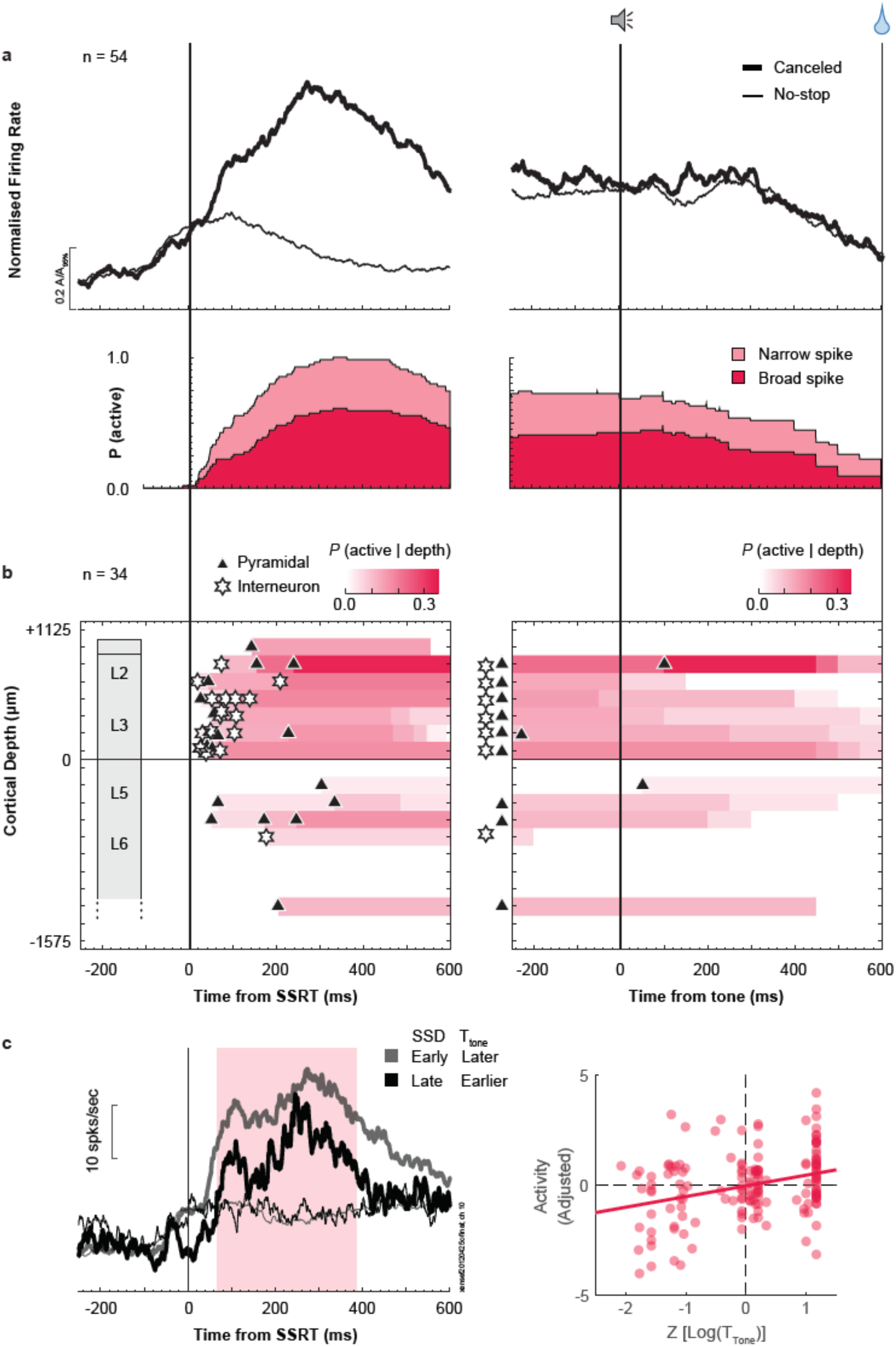
Time-depth organization of Goal Maintenance neuron spiking in SEF. **a,** Normalized population response of neurons with prolonged facilitation in discharge rate on successful canceled (thick) relative to latency-matched no stop-signal (thin) trials for early SSD. **b,** Recruitment of this signal through time relative to SSRT (left) and auditory feedback tone (right), with dark and light shades representing the recruitment of broad-spiking (spike width ≥ 250 µs) and narrow-spiking (< 250 µs) neurons. Recruitment on SSRT-aligned activity (left panel) is defined as the difference between canceled and no stop-signal trials. Recruitment on tone-aligned activity (right panel) is defined as the activity on canceled trials relative to the baseline. Modulations starting 300ms after the tone are not shown. **c,** Time-depth plot showing latency and proportion of recruited neurons through time at each depth from perpendicular penetrations. Symbols mark beginning of modulation for broad-spiking neurons (black triangles) and narrow-spiking neurons (white stars). Color map indicates the percentage of neurons relative to the overall sampling density (**Supplementary Figure 1a**) producing this signal through time at each depth. Dashed horizontal line marks L3-L5 boundary. The lower boundary of L6 is not discrete. **d,** Left panel compares response of a representative neuron on successfully canceled (thick) relative to latency-matched no stop-signal (thin) trials for early (lighter) and later (darker) SSD. Shaded area represents significant difference in discharge rate between the two conditions. Right panel plots relationship between discharge rate in the sampling interval and feedback tone time. Along the ordinate scale is plotted the spiking rate, adjusted for neuron-specific variations. Along the abscissa scale is plotted the normalized feedback time in logarithmic scale (z-scale). In all, 162 points (54 neurons) are plotted. Each point plots the average spike-density and associated Log (feedback time) in one of 3 bins corresponding to early-, mid-, or late-SSD, for each neuron. Variation of spiking rate was best predicted by the time of the feedback time (highlighted by best-fit line; **Supplementary Table 2**).

These neurons were distinguished from Conflict neurons by the more prolonged facilitation following SSRT (**Supplementary Figure 1b, c**). The peak recruitment of these neurons (∼300 ms) followed that of the neurons monitoring conflict (∼110 ms) and the suppression of the Event Timing neurons (∼170 ms). Compared to Conflict neurons, the phasic facilitation was followed by sustained activity until ∼300 ms after the feedback tone in a significantly higher proportion of these neurons (*X*^2^ (1, *N* = 129) = 27.3, p < 10^-5^) (**Fig 4a**). This modulation at tone presentation was also observed on no stop-signal trials. The variation in the magnitude of the phasic modulation was best described by the log-transformed duration until the feedback tone on canceled trials (**Fig 3d**) (t(152) = 3.53, p = 5.6×10^-4^), with strong evidence against non-time-based models (Δ BIC > 3.0) and weak evidence against other time-based models (ΔBIC < 1) (**Supplementary Table 2**).

In a large proportion of these neurons, the phasic response on canceled trials after SSRT was followed by a sustained elevated discharge rate that was interrupted after the tone. This sustained activity was also observed on no-stop trials. Consistent with the indirect contribution of SEF to saccade initiation, the termination of this modulation was unrelated to when monkeys stopped fixating on the stop-signal (on canceled trials) or the target (on no-stop trials), ruling out this signal as one directly involved in maintaining fixation (**Supplementary Figure 5c**). Furthermore, when the feedback tone cued upcoming reward, the activity was suppressed; when the tone cued failure, activity increased (**Supplementary Figure 5d**). Accordingly, by representing both time and valence of the feedback tone, a significant proportion of these neurons also signaled Loss as described previously ^16^ (*X*^2^ (3, *N* = 575) = 19.43, p = 2.2×10^-4^; **Supplementary Table 3**). Based on the observation that this activity was sustained until the tone, which signaled when gaze could be shifted, and previous findings identifying SEF signals with working memory ^10, 11^, we conjecture that these neurons sustain saccade inhibition to earn reward. Hence, we refer to these neurons as Goal Maintenance neurons.

Goal Maintenance neurons were found in all penetrations but more commonly at certain sites (*X*^2^ (4, *N* = 575) > 39.3, p < 10^-5^). One third (18/54) were narrow-spiking, a proportion exceeding chance sampling (*X*^2^ (1, *N* = 575) = 7.29, p = 0.0069). The laminar distribution of Goal Maintenance neurons (**Fig. 4c**) was significantly different from the laminar sampling distribution (*X*^2^ (4, *N* = 293) = 11.24, p = 0.024) (**Supplementary Table 1b**). These neurons were found significantly more often in L2/3 relative to L5/6 (*X*^2^ (1, *N* = 293) = 10.37, *p* = 1.3×10^-4^). Their laminar distribution was also significantly different from that of Conflict neurons (*X*^2^ (1, *N* = 70) = 11.54, *p* = 6.8×10^-4^) and of Event Timing neurons (*X*^2^ (1, *N* = 83) = 5.49, *p* = 0.019). Those in L2/3 modulated significantly earlier than those in L5/6 (L2/3 ∼85 ± 64 ms (mean ± SD), L5/6 ∼ 193 ± 101; *t*-test, t(32) = −3.63, p = 9.9×10^-4^).

In summary, consistent with previous studies ^10, 11^, neurons in SEF produce activity sufficient to enable a working memory representation of the goal of saccade inhibition through time. The new results show that these neurons are most common in L2/3 and a relatively higher proportion have narrow spikes. Thus, at least some of these neurons can be inhibitory interneurons.

### Countermanding N2

To determine whether macaque monkeys produce ERP components associated with response inhibition homologous to humans ^5^, we simultaneously sampled EEG from an electrode located over the medial frontal cortex (Fz in 10-20 system) while recording neural spikes in SEF (**Fig. 5a**). To eliminate components associated with visual responses and motor preparation and to isolate signals associated with response inhibition, we measured the difference in polarization on canceled trials and latency-matched no stop-signal trials for each SSD (**Fig. 5b**). Homologous to humans, we observed an enhanced N2/P3 sequence with successful stopping.

**Fig. 5.**
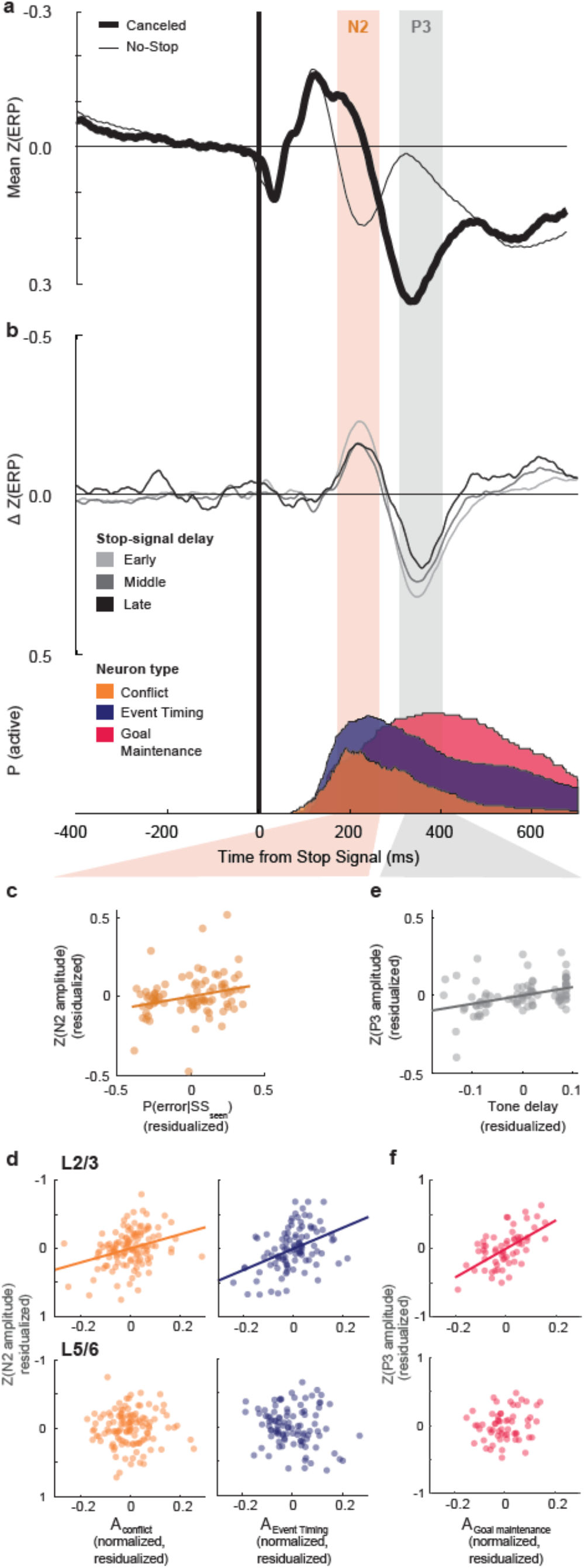
Event-related potentials for successful response inhibition. **a,** Grand average normalized EEG (z-transformed) on successful canceled (thick) relative to latency-matched no stop-signal (thin) trials for early SSD. **b,** the difference function highlights the N2 and P3 components, eliminating the effect of response stimulus-evoked ERP common to both canceled and no stop-signal trials. The shaded regions correspond to a ±50 ms sampling window around peak of N2 (orange) and P3 (gray) used for ERP amplitude calculation for **c**. **c,** Relationship between N2 amplitude and P(error | SS_seen_). Along the ordinate scale, the normalized ERP amplitude is plotted, adjusting for session-specific variations. Along the abscissa scale the normalized P(error | SS_seen_) is plotted (**Supplementary** Fig 2c). In all, 87 points (29 sessions) are plotted. Each point plots the average N2 and the associated P(error | SS_seen_) in one of 3 bins corresponding to early-, mid-, or late-SSD, for each session. P(error | SS_seen_) is the best parameter that described variations in N2 (highlighted by best-fit line). **d,** Relationship between P3 amplitude and the time of feedback relative to stop-signal. Along the ordinate scale is plotted the normalized ERP amplitude (z-scale), adjusted for session-specific variations in amplitude. Along the abscissa scale is plotted the normalized feedback time in logarithmic scale (z-scale). In all, 87 points (29 sessions) are plotted with each point plotting the average spike-density and associated Log (feedback time) in one of 3 bins corresponding to early-, mid-, or late-SSD, for each neuron. Variation of P3 amplitude was best predicted by the time of the feedback time (highlighted by best-fit line; **Supplementary Table 2**). **e,** Relationship between laminar neuronal discharge rate and N2. From sessions with perpendicular penetrations, relationship between ERP amplitude and spike rate for Conflict neurons (*A*_Conflict_), Event Timing neurons (*A*_Event Timing_), recorded in L2/3 (top) and L5/6 (bottom). Partial regression plots are obtained by plotting on the ordinate scale, according to EEG convention, the residual from fixed-effects-adjusted ERP amplitude controlling for activity in the opposite layer. Along the abscissa scale is plotted the residual fixed-effects-adjusted neuronal discharge rate in the identified layer controlling for the activity in the opposite layer and stop-signal delay. Each point plots the average EEG voltage and associated spiking rate in one of 20 bins with equal numbers of trials per session. Only sessions with neurons in both L2/3 and L5/6 are included. A total of 120 points (from 6 session) are plotted for Conflict Neurons (left), and 100 points (5 sessions) are plotted for Event Timing neurons (right). The relationship between N2 and other neurons not reported in this study and Goal Maintenance neurons are shown in **Supplementary Fig 7a**. Variations in N2 amplitude was predicted by variation of spiking rate of Conflict and Event Timing neurons in L2/3 (highlighted by best-fit line) but not in L5/6. **f,** Relationship between laminar neuronal discharge rate and P3. From sessions with perpendicular penetrations, relationship between ERP amplitude and spike rate for Goal Maintenance neurons (*A*_Goal Maintenance_), recorded in L2/3 (top) and L5/6 (bottom). Partial regression plots are obtained by plotting on the ordinate scale, according to EEG convention, the residual from fixed-effects-adjusted ERP amplitude controlling for activity in the opposite layer and stop-signal delay. Similar conventions to panel **e**. Only sessions with neurons in both L2/3 and L5/6 are included. A total of 60 points (from 3 sessions) are plotted for Goal Maintenance neurons. The relationship between P3 and other neuron classes are shown in **Supplementary Fig 7c**. Variations in P3 amplitude was predicted by variation of spiking rate of Goal Maintenance neurons in L2/3 (highlighted by best-fit line) but not in L5/6.

The N2 began ∼150 ms and peaked 222 ± 17 ms after the stop-signal, well after the visual ERP polarization (**Supplementary Fig 6a**). The N2 was observed after SSRT, too late to be a direct index of reactive response inhibition. Furthermore, the variability in the N2 peak time across sessions was significantly less when aligned on stop-signal appearance than on SSRT, further dissociating the N2 from reactive inhibition (F-test for variances, F(28,28) = 0.29, p = 0.0018) (**Supplementary Fig 6c**). N2 amplitude varied most with P(error | SS_seen_) (Δ BIC > 3.0 against all competing models), with the largest negativity during the earliest SSD associated with the lowest error likelihood (t(85) = 2.42, p = 0.0178) (**Fig 5c, Supplementary Table 2**). In fact, no other competing model explained the variation in N2 amplitude. This outcome adds to the inconsistent and inconclusive evidence for the N2 association with conflict monitoring and response inhibition ^5^.

We now describe relationships between neural spiking and the N2. **Figure 5b** illustrates the temporal relationship between the ERP and the recruitment of the three classes of neurons described above. The N2 coincided with the peak recruitment of Conflict and of Event Timing neurons. The relationship between neural events in SEF and the voltages measured on the cranium above SEF is both biophysical and statistical. The cranial voltage produced by synaptic currents associated with a given spike must follow Maxwell’s equations as applied to the brain and head, regardless of the timing of the different events. Hence, we counted the spikes of the three classes of neurons separately in L2/3 and in L5/6 during a 100 ms window centered on the peak of the ERP. We devised multiple linear regression models with activity in upper layers (L2/3) and lower layers (L5/6) of each neuron class as predictors. Only successfully canceled trials were included in this analysis. We found that variation in the polarization of the N2 is not associated with the phasic spiking of Goal Maintenance neurons (L2/3: t(57) = −1.28, p = 0.21; L5/6: t(57) = 0.60, p = 0.52) (**Supplementary Figure 7a**) but was predicted by the spiking activity in L2/3 but not in L5/6 of Conflict (L2/3: t(117) = −3.6, p = 4.7×10^-4^; L5/6: t(117) = 0.046, p = 0.96) and of Event Timing neurons (L2/3: t(97) = −4.60, p = 1.3×10^-5^; L5/6: t(97) = 1.67, p = 0.097) (**Fig 5d**). When the discharge rate of these L2/3 neurons was higher, the N2 exhibited a stronger negativity. Interestingly, N2 polarization was also predicted by the spiking activity in L2/3 but not in L5/6 of other neurons that were not modulated on canceled trials and so were not described in this manuscript (L2/3: t(317) = −2.51, p = 0.012; L5/6: t(317) = −1.60, p = 0.11).

Similar results were obtained when controlling for the variation of ERP polarization and spike rate across different SSDs (not shown) and when measuring the difference in spiking and ERP between canceled and matched no stop-signal trials (**Supplementary Figure 7b**).

### Countermanding P3

The N2 was followed by a robust P3 (**Fig 5a, b**) beginning ∼300 ms and peaking 358 ± 17 ms after the stop-signal, homologous to the human P3 ^5^. The peak polarization time was better synchronized on the stop-signal than on SSRT (F(28,28) = 0.44, p = 0.0345) (**Supplementary Fig 6c**). P3 polarization varied most with the log-transformed time of the feedback tone on canceled trials (Δ BIC > 4.0 against competing models) with weak support against other time-based models (Δ BIC < 1.30) (**Fig 5e, Supplementary Table 2**). P3 polarization increased with time until feedback (t(85) = 3.72, p = 3.5×10^-4^). The conclusions of these results do not differ if the analyses are performed on the raw EEG polarization in these intervals.

Peak P3 polarization coincided with the peak recruitment of Goal Maintenance neurons, while the recruitment of Conflict and Event Timing neurons was decaying (**Fig 5b**). Accordingly, variation in P3 polarization was predicted by the spiking activity of Goal Maintenance neurons in L2/3 but not L5/6 (L2/3: t(57) = 5.46, p = 1.1×10^-6^; L5/6: t(57) = 1.47, p = 0.15) (**Fig. 5f**). Higher spike rates are associated with greater P3 positivity. P3 amplitude was not associated with the spiking of Conflict (L2/3: t(97) = 0.44, p = 0.66; L5/6: t(97) = −0.49, p = 0.62), Event Timing (L2/3: t(117) = −1.19, p = 0.24; L5/6: t(117) = −0.78, p = 0.44), or unmodulated neurons (L2/3: t(317) = −1.11, p = 0.27; L5/6: t(317) = 0.054, p = 0.96) (**Supplementary Figure 7c**). Similar results were obtained when SSD was controlled for (not shown) and when measuring the difference in spiking and ERP between canceled and matched no stop-signal trials (**Supplementary Figure 7d**).

## DISCUSSION

These results offer important, new insights into the cortical microcircuitry supporting executive control in primates. Model-based analysis of the latency, temporal dynamics, and variation in strength of neural spiking across the neuron sample revealed functionally distinct and theoretically important classes of neurons with particular biophysical and laminar properties. Moreover, a bridge between these neurophysiological findings and human electrophysiology was established through the specific associations observed between the N2 and P3 ERP observed in response inhibition tasks and classes of neurons in particular cortical layers. The novelty and importance of these findings is amplified by their complementarity with our previous description of the laminar organization of error and reward processing in SEF ^16^. Based on the new results, we will discuss how SEF can contribute to conflict monitoring, time estimation, and goal maintenance. Coupled with extensive knowledge about connectivity of SEF ^42, 43, 44^, this new information about the laminar distribution of neurons signaling response conflict, event timing, and maintaining goals suggest several specific hypotheses and research questions about how SEF and associated structures accomplish response inhibition and executive control (**Fig. 6**). Also, complementing our earlier description of the source of the ERN ^16^, we now report a macaque homolog of the N2/P3 ERP components associated with response inhibition. The new results demonstrate one cortical source of these ERP components.

**Fig 6.**
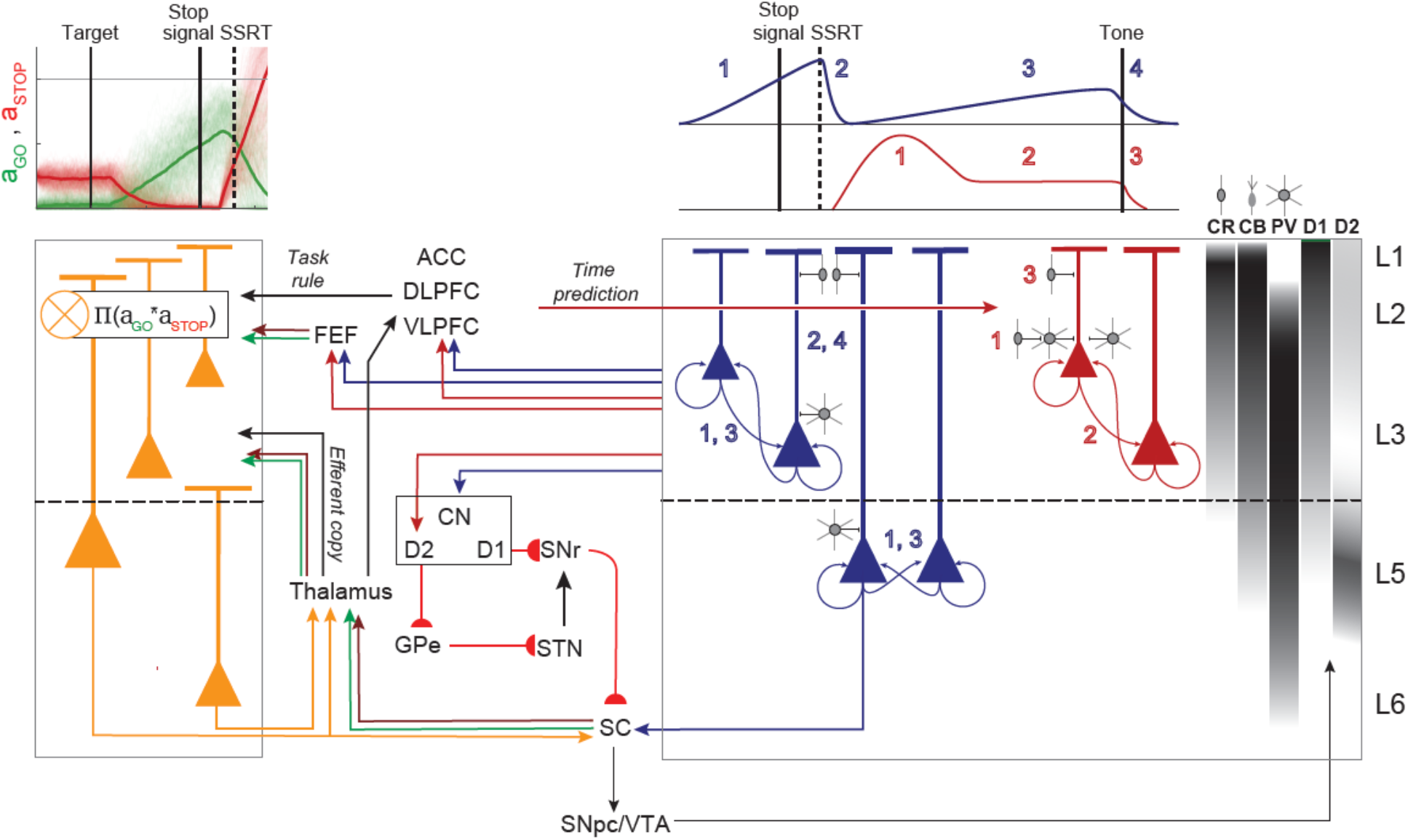
Extrinsic and intrinsic circuitry for executive control. The laminar distribution observed for Conflict (orange), Event Timing (dark blue), and Goal Maintenance (dark red) are summarized with selected anatomical connections based on published studies. Sampled neurons were likely broad spike pyramidal and narrow spike, possibly inhibitory neurons. The laminar densities of calretinin (CR), calbindin (CB), and parvalbumin (PV) neurons observed and of D1 and D2 receptors are indicated on the far right. **Left**, Conflict signal can arise in SEF through afferents from frontal eye field (FEF). SEF can receive coincident inputs from Fixation neurons (STOP) and Movement neurons (GO) in FEF, directly, or in SC, indirectly, via thalamus, terminating in L2/3. These inputs are integrated within the synapses of L2/3 and L5 Conflict neurons. Intracortical processing produces later activation of Conflict neurons in L6 which can relay this signal to the Thalamus. **Right**, Top: Schematic of the activity profile for Goal Maintenance and Event Timing neurons in distinct phases indicated by the number. We conjecture that Goal Maintenance neurons, mainly located in L2/3, suppress unwanted movement through push-pull basal ganglia circuitry with pyramidal neurons directly projecting to the indirect pathway (D2) and inhibitory neurons, inhibiting pyramidal neurons that can project to the direct (D1) pathway. The gray symbol indicates that these neurons are distinct from those reported in this study. Input from dorsolateral prefrontal cortex (DLPFC) and anterior cingulate cortex (ACC), terminating in L2/3 can inform SEF of the anticipated reward association based on the experienced stop-signal delay contingent on successful stopping. Dopamine (DA) neuron projections in L2/3 from the SNpc and VTA can also relay this information. These inputs can result in the phasic response in Goal Maintenance neurons (phase 1, red). Following the phasic response, activity can remain elevated via recurrent connections and balance of excitation and inhibition (phase 2, red). The auditory feedback tone, integrated with the task rule from DLPFC cues the termination of operant control on behavior, resulting in the inhibition of pyramidal and inter-neurons by CR and CB neurons. This results in the termination of the sustained activity (phase 3). Event Timing neurons can receive input from DLPFC and ACC terminating in L2/3 informing neurons in L2/3 and L5 about an upcoming event. Ramping results from recurrent connections (1, dark blue). SEF can receive information about stop-signal appearance and successful stopping from ventrolateral prefrontal cortex (VLPFC) and DLPFC and Conflict neurons within the microcircuitry. This information can suppress the ramping activity via inhibitory connections by direct inhibitory connections onto Event Timing neurons (phase 2, dark blue). This resets these neurons for the next phase of ramping (phase 3, dark blue) which is terminated by the appearance of the feedback tone (4). The activity of Event Timing neurons can project to the caudate nucleus to inform the fronto-striatal reinforcement learning loop about the experienced timing of the event. Further details in text.

### Conflict

One class of SEF neuron was characterized by a pronounced facilitation after the stop-signal when saccades were inhibited. The modulation followed SSRT and scaled with P(error). These neurons were predominantly broad spiking and found in all layers. We hypothesize that these neurons signal response conflict ^15, 39^ defined as co-activation of mutually incompatible response processes ^31^. Previous research has characterized the neural mechanism of saccade countermanding ^27, 45^. On canceled trials, gaze-shifting and gaze-holding neurons in the frontal eye field (FEF) and superior colliculus (SC) are co-active in a dynamically unstable manner that varies with P(error) precisely because these are the neurons producing the performance. In the interactive race model ^29, 30^, the multiplicative conflict between GO and STOP accumulator units scales with P(error) (**Supplementary Figure 2a**) and can be used to adjust interactive race parameters to accomplish post-stopping slowing ^39^. Thus, these neurons signal a quantity central to theories of executive control. Furthermore, different neurons in SEF signal conflict, error, and reward, highlighting the possible independence of these executive control signals.

Further evidence dissociating conflict, reward, and error signals is offered by comparing our results with those of a recent investigation of the nigrostriatal dopamine system of monkeys performing saccade countermanding ^46^. Dopamine (DA) neurons concentrated in the dorsolateral substantia nigra exhibited a pattern of activity that paralleled the conflict neurons in SEF. The DA neurons produced a brisk response to the stop-signal that was stronger when saccades were canceled in either direction. This observation is consistent with reports that besides responding to rewarding events, dopamine neurons respond to salient signals, such as a stop-signal. Unlike movement neurons in FEF ^27^ and SC ^45^ but like SEF, nearly all DA neurons modulate after SSRT. Moreover, the modulation of DA neurons scaled with P(error) just like the SEF neurons.

The striking parallels between SEF and SNpc modulation patterns invites consideration of cause and effect. SEF is innervated by DA neurons in Substantia Nigra pas compacta (SNpc) ^47^, and SNpc DA neurons modulated significantly earlier than did the SEF conflict neurons (**Supplementary Figure 8**). However, because of the very slow conduction of DA axons ^48, 49, 50, 51^, we estimate that the spike conduction time from SNpc to SEF is ∼100 ms (**Supplementary Figure 8**). Consequently, the estimated arrival times of DA spikes in SEF were not significantly different from the modulation times of the conflict neurons (**Supplementary Figure 8**). The influence of DA in SEF is slowed further by the well-known second-messenger delay of influence. Therefore, we infer that the SEF conflict modulation cannot be caused by DA inputs. However, because axon terminals from SEF are rare in SNpc ^42, 44^, SEF neurons are unlikely to cause directly the modulation of the SNpc DA neurons. Instead, other investigators have shown that the phasic DA activation is delivered by the SC ^52^. Through the conflict neurons in L5, SEF can influence SC directly ^42^. Curiously, though, the modulation specifically after SSRT scaling with P(error) has not been observed in SC ^45^.

Theories of DA function can facilitate understanding the putative conflict signal in SEF. From the reinforcement perspective, the phasic DA signal may act as an immediate eligibility trace broadcast to SEF and other regions to associate reinforcement with successful cancelation to the infrequent stop-signal. Such eligibility traces must be salient to be useful. The reinforcement perspective suggests an alternative to the conflict interpretation. The imbalance between gaze-holding and gaze-shifting arising on canceled trials increases with the progressive commitment from gaze-holding to gaze-shifting through time. Consequently, as the likelihood of unsuccessful response inhibition increases, the surprise of successful response cancelation increases. We observed a divergence in the values of P(error)—which is necessarily proportional to the product of the activation of GO and STOP units—and P(error | SSseen)—which is a proxy of error likelihood learned through experience with the task—at longer SSDs. Others have described SEF neural signals in terms of surprise ^12^. Thus, the modulation after SSRT scaling with P(error) may just be another element of the reinforcement learning needed to perform this task. Further research is needed to resolve the conflict and surprise hypotheses.

Conflict neurons were found in all layers. To signal conflict, SEF can be informed about the dynamic state of gaze-shifting and gaze-holding through inputs from FEF and oculomotor thalamic nuclei. To signal surprise, SEF can be informed about saccade production from the thalamus ^53^ and task rules from DLPFC and ventrolateral prefrontal cortex (VLPFC) ^54^. Based on previous conjectures ^6^ and recent biophysical modelling ^55^ we hypothesize that the integration of information producing the modulation of these neurons is derived through synaptic processes in L2/3. However, if this is so, and if the apical dendrites of L6 pyramidal neurons in SEF do not extend into L2/3, then this conflict signal can arise in L6 through translaminar interactions. The observation that conflict arises later in L6 is consistent with this supposition. Another implication of the hypothesis that conflict in L6 is derived from that in L2/3 is that the L6 feedback to thalamus will be delayed relative to the gaze-holding and gaze-shifting signals conveyed from the SC.

### Time estimation and goal maintenance

The interpretation of the other two classes of neurons that we found is framed by motivation more than reinforcement. To earn reward, monkeys must hold gaze for an extended period, which requires preventing blinks that would interrupt the camera-based eye tracker. This entails learning and possibly exploiting any regularities in the timing of task events ^33, 40^. A contribution of SEF and nearby areas in action timing and explicit time production tasks has been demonstrated ^13, 14^. We extend that description to this stop-signal task.

A distinct group of SEF neurons produced ramping activity before saccades, which decayed after the gaze shift. But, when the saccade was countermanded, the ramping was interrupted by pronounced suppression. A previous description of these neurons recognized that the modulation on canceled trials arose too late to contribute to reactive inhibition but offered no explanation for these neurons ^7^. The new task design used here exposed a second period of ramping before the feedback tone on ∼30% of these neurons. This monotonically rising activity reached different levels for different interval durations ranging from ∼1000 to 1400 ms after SSRT on canceled trials. Our discovery of an association between spiking rate and the log-transformed duration of the preceding interval motivates a more integrated interpretation framed by a body of research on time keeping ^37, 41^^56 57^.

We interpret the ramping activity as representing the timing of task events. Spiking rate increases as the learned time of an event like the stop-signal approaches. Strong suppression after the event resets a proportion of these neurons to ramp until the next event, i.e., the feedback tone. The stop-signal and feedback tone events differ in two ways. First, they differ in predictability, for the stop-signal only occurs on a proportion of trials while feedback tone is not. Second, they differ in the action required following the event, for the stop-signal announces a prolonged period of fixation in which blinks must also be withheld while the tone announces the release of control over behavior.

Recent work has shown that different neurons in the basal forebrain signal timing of events depending on surprise, salience, and uncertainty ^37^. We found similar differences in SEF. We conjecture that those neurons with ramping activity before both SSRT and the feedback tone encode the timing of expected salient events regardless of certainty or expected action. In contrast, the neurons with only ramping activity before successful stopping encode events that are less certain in occurrence or consequence. These differences were reinforced by the distribution of the neurons across the cortical layers. While Event Timing neurons were found in all layers, those that encoded timing regardless of predictability or action were most common in L3 and L5 with broad spikes consistent with pyramidal projection neurons.

This laminar differentiation demonstrates that the timing of different types of events can engage different circuits mediated by different layer-specific extrinsic connections. The timing signal can be sent via cortico-cortical connections to other cortical areas to influence motor, cognitive, and limbic processes. Further research is needed to clarify this projection. Also, these neurons can contribute to fronto-striatal pathways to learn and update the temporal structure of the task ^57, 58, 59^. Axon terminals from SEF are dense in the caudate nucleus ^43^, arising from pyramidal neurons in L3 and L5 ^60, 61, 62^. In fact, neurons with this pattern of modulation have been described in a recent investigation of the caudate nucleus of monkeys performing saccade countermanding ^46^. Our finding that the suppression in the caudate nucleus occurred significantly later after SSRT than that of Event Timing neurons in SEF (**Supplementary Figure 8**) suggests a primary role of the cortex in this signaling.

The rapid suppression of the ramping activity after SSRT merits consideration. One source can be intracortical inhibition from the narrow-spike, putative PV neurons that we observed. Another source can be the very small CB and CR neurons concentrated in L2/3 that are innervated by DLPFC and selectively inhibit pyramidal neurons ^63^, although our methods are unlikely to sample spikes from them. We note that although SEF is an agranular structure with weak interlaminar inhibitory connections ^21^, CR neurons in L2/3 can potently inhibit L5 neurons through specialized projections on the apical dendrites ^64^. This inhibition must be informed about the presence of the stop-signal and the cancelation of the saccade. We observe that such a signal is available in the conflict neurons. However, the suppression of Event Timing neurons occurred significantly earlier than the facilitation of the conflict neurons. Further research can resolve these cortical interactions.

The Event Timing neurons that represent the duration of a preceding interval can support the patterns of modulation observed in the final class of neuron we found. The third class of neuron produced a phasic response after SSRT on canceled trials that scaled with the duration of the upcoming interval until the feedback tone. Recall that on canceled trials the interval from target presentation until tone presentation was of fixed duration, making it progressively shorter after progressively longer SSD. Such phasic responses have previously been observed when the timing of events followed discrete predictable durations ^65^ similar to the time of feedback tone in our task following successful stopping. This phasic representation of the time was followed by sustained spiking until the tone. Note that by design, when the tone sounded, monkeys could shift gaze before receiving the fluid reward. We propose that these neurons can be identified with the operation of goal maintenance, which is necessary in canceled trials to prevent blinking or gaze shifts before the tone. This inference is consistent with an interpretation of the original theory of response inhibition ^26^ and supported by previous evidence linking SEF to working memory ^10, 11^ and working memory to time representation ^66, 67^. We have obtained further evidence for this interpretation in ongoing experiments with two other monkeys performing the same saccade countermanding task but with the requirement to maintain fixation on the stop-signal until the fluid reward is delivered. Goal maintenance neurons have been observed, but they continue spiking after the tone until the fluid reward when operant control over behavior is released (data not shown).

Goal maintenance neurons were mainly found in L2/3. Inputs to these neurons from DLPFC, VLPFC, and ACC can signal task rules and the expected time of the secondary reinforcer when executive control can be released. Dopaminergic release in SEF from VTA where similar time-predicting signals are observed ^65^ can enhance these influences through higher density of D1 relative to D2 receptors in L2/3 ^68, 69^. The sustained discharge can be maintained through recurrent activation within SEF and between other structures ^11, 70^. Also, many goal maintenance neurons had narrow spikes, consistent with PV inhibitory neurons, which can balance excitation and inhibition necessary for the maintenance of persistent activity in recurrent networks ^71,72, 73, 74^.

We hypothesize that pyramidal Goal Maintenance neurons can encourage the suppression of movements through projections to the indirect pathway D2 neurons in the striatum ^60, 61, 62^. Inhibitory Goal Maintenance neurons, on the other hand, can inhibit the D1 direct (action-promoting) pathway and the frontal eye field to suppress movement. As PV neurons in primates do not have extrinsic connections, we propose that this can be mediated by the inhibition of other excitatory neurons (unidentified neurons and possibly Gain neurons identified in ^16^) that send projections to these motor structures (gray neurons). Therefore, Goal Maintenance neurons can achieve their role by altering the balance in the push-pull mechanism mediated by the direct (D1) and indirect (D2) pathways. This function is consistent with the observation that many of these neurons also exhibit higher activity on unrewarded trials that, as previously described, influences post-error adjustments in RT in the next trial ^16^. Also consistent with this hypothesis, neurons with facilitated activity after SSRT were described in an investigation of the caudate nucleus of monkeys performing saccade countermanding ^46^. The facilitation in the caudate nucleus coincided with that measured in SEF (**Supplementary Figure 8**). The parallel between SEF and the striatum in patterns of modulation associated with proactive but not reactive inhibition are surprisingly, but satisfyingly, clear.

### Origin of Countermanding N2/P3

We showed that macaque monkeys exhibit a N2/P3 ERP complex homologous to that observed in humans ^5^. We discovered that variation in N2 and P3 polarization was predicted by spiking of specific, different neuron classes in L2/3 and not L5/6. These findings extend and parallel our previous demonstration that SEF contributes to the error-related negativity (ERN) ^16^. We found that variations in error-related spiking in L2/3 but not in L5/6 predicted variation of EEG polarization across both error and correct trials. Because action potentials are not large or sustained enough to produce event-related potentials, we surmise that this neural spiking coincides with coherent current flow strong enough to produce in the ERN ^55^. These new results show synaptic activity in L2/3 of SEF contributes to the N2/P3 complex.

Disagreement persists about what the frontal N2 indexes ^75, 76^. We found that the amplitude of the macaque homologue of the N2 during the stop-signal task varied most with the likelihood of error associated with experienced SSDs and not conflict and SSD as previously suggested ^5, 77^. Further, we demonstrate that the spiking of different classes of neurons in L2/3 (but not L5/6) predicted the magnitude of the N2. Specifically, N2 magnitude was unrelated to spiking of Goal Maintenance neurons but co-varied with spiking of Conflict and Event Timing neurons in addition to the spiking of other neurons that did not modulate around the time of successful stopping. Recognizing that the N2 manifests the influence of different processes occurring in functionally distinct neurons can explain the disagreement about the nature of this ERP component.

Likewise, the macaque homologue of the P3 component in this task resembled that reported in humans ^5^. Consistent with previous reports of P3 indexing expectation and temporal aspects of behavior ^75^, we found that P3 amplitude co-varied most with the expected time of the feedback tone. Reinforcing this interpretation, we found that P3 amplitude was predicted by the spiking of Goal Maintenance but not Conflict or Event Timing neurons. Therefore, we surmise that the P3 expressed in our experimental design indexes temporal prediction underlying goal maintenance. Overall, these results demonstrate that N2 and P3 index distinct processes mediated by the activity of different populations of neurons.

## Conclusion

Pioneering insights into the microcircuitry and mechanisms of primary visual cortex began by describing the properties of neurons in different layers ^78^. The present results complete the first catalogue for an agranular frontal lobe area. Contrasts with primary sensory areas will reveal the degree of computational uniformity across cortical areas. Being a source contributing to ERPs indexing performance monitoring and executive control, details about laminar processing in SEF will offer unprecedented insights into the microcircuitry of executive control. These results validate the tractability of formulating neural mechanism models of performance monitoring and executive control, especially when constrained by formal ^26^, algorithmic ^29, 30^, and spiking network ^79^ models of performance of a task with clear clinical relevance ^80^.

## METHODS

### Animal care and surgical procedures

Data was collected from one male bonnet macaque (Eu, *Macaca Radiata*, 8.8kg) and one female rhesus macaque (X, *Macaca Mulatta*, 6.0kg) performing a countermanding task ^20, 24^. All procedures were approved by the Vanderbilt Institutional Animal Care and Use Committee in accordance with the United States Department of Agriculture and Public Health Service Policy on Humane Care and Use of Laboratory Animals. Surgical details have been described previously ^81^. Briefly, magnetic resonance images (MRIs) were acquired with a Philips Intera Achieva 3T scanner using SENSE Flex-S surface coils placed above or below the animal’s head. T1-weighted gradient-echo structural images were obtained with a 3D turbo field echo anatomical sequence (TR = 8.729 ms; 130 slices, 0.70 mm thickness). These images were used to ensure Cilux recording chambers were placed in the correct area. Chambers were implanted normal to the cortex (Monkey Eu: 17°; Monkey X: 9°; relative to stereotaxic vertical) centered on midline, 30mm (Monkey Eu) and 28mm (Monkey X) anterior to the interaural line.

### Acquiring EEG

EEG was recorded from the cranial surface with electrodes located over medial frontal cortex. Electrodes were referenced to linked ears using ear-clip electrodes (Electro-Cap International). The EEG from each electrode was amplified with a high-input impedance head stage (Plexon) and bandpass filtered between 0.7 and 170 Hz. Trials with blinks within 200ms before or after the analysis interval were removed.

### Cortical mapping and electrode placement

Chambers implanted over the medial frontal cortex were mapped using tungsten microelectrodes (2-4 MΩ, FHC, Bowdoin, ME) to apply 200ms trains of biphasic micro-stimulation (333 Hz, 200 µs pulse width). The SEF was identified as the area from which saccades could be elicited using < 50 µA of current ^82, 83^. In both monkeys, the SEF chamber was placed over the left hemisphere. The dorsomedial location of the SEF makes it readily accessible for linear electrode array recordings across all cortical layers. A total of five penetrations were made into the cortex—two in monkey Eu, three in monkey X. Three of these penetration locations were perpendicular to the cortex. In monkey Eu, the perpendicular penetrations sampled activity at site P1, located 5 mm lateral to the midline and 31 mm anterior to the interaural line. In monkey X, the perpendicular penetrations sampled activity at site P2 and P3, located 5 mm lateral to the midline and 29 and 30 mm anterior to the interaural line, respectively. However, during the mapping of the bank of the cortical medial wall, we noted both monkeys had chambers place ∼1 mm to the right respective to the midline of the brain. This was confirmed through co-registered CT/MRI data. Subsequently, the stereotaxic estimate placed the electrodes at 4 mm lateral to the cortical midline opposed to the skull-based stereotaxic midline.

### Acquiring neural spiking

Spiking activity and local field potentials were recorded using a 24-channel Plexon U-probe with 150 µm between contacts, allowing sampling from all layers. The U-probes were 100 mm in length with 30 mm reinforced tubing, 210 µm probe diameter, 30° tip angle, with 500 µm between the tip and first contact. Contacts were referenced to the probe shaft and grounded to the headpost. We used custom built guide tubes consisting of 26-gauge polyether ether ketone (PEEK) tubing (Plastics One, Roanoke, VA) cut to length and glued into 19-gauge stainless steel hypodermic tubing (Small Parts Inc., Logansport, IN). This tubing had been cut to length, deburred, and polished so that they effectively support the U-probes as they penetrated dura and entered cortex. The stainless-steel guide tube provided mechanical support, while the PEEK tubing electrically insulated the shaft of the U-probe, and provided an inert, low-friction interface that aided in loading and penetration.

Microdrive adapters were fit to recording chambers with <400 µm of tolerance and locked in place at a single radial orientation (Crist Instruments, Hagerstown, MD). After setting up hydraulic microdrives (FHC, Bowdoin, ME) on these adapters, pivot points were locked in place by means of a custom mechanical clamp. Neither guide tubes nor U-probes were removed from the microdrives once recording commenced within a single monkey. These methods ensured that we were able to sample neural activity from precisely the same location relative to the chamber on repeated sessions.

Electrophysiology data were processed with unity-gain high-input impedance head stages (HST/32o25-36P-TR, Plexon). Spiking data were bandpass filtered between 100 Hz and 8 kHz and amplified 1000 times with a Plexon preamplifier, filtered in software with a 250 Hz high-pass filter and amplified an additional 32,000 times. Waveforms were digitized at 40 kHz from −200 to 1200 µs relative to voltage threshold crossings. Thresholds were typically set at 3.5 standard deviations from the mean. All data were streamed to a single data acquisition system (MAP, Plexon, Dallas, TX). Time stamps of trial events were recorded at 500 Hz. Single units were sorted online using a software window discriminator and refined offline using principal components analysis implemented in Plexon offline sorter.

### Cortical depth and layer assignment

The retrospective depth of the electrode array relative to grey matter was assessed through the alignment of several physiological measures. Firstly, the pulse artifact was observed on a superficial channel which indicated where the electrode was in contact with either the dura mater or epidural saline in the recording chamber; these pulsated visibly in synchronization with the heartbeat. Secondly, a marked increase of power in the gamma frequency range (40-80Hz) was observed at several electrode contacts, across all sessions. Previous literature has demonstrated elevated gamma power in superficial and middle layers relative to deeper layers ^84, 85^. Thirdly, an automated depth alignment procedure was employed which maximized the similarity of CSD profiles evoked by passive visual stimulation between sessions ^20^.

Further support for the laminar assignments was provided by an analysis of the depths of SEF layers measured in histological sections visualized with Nissl, neuronal nuclear antigen (NeuN), Gallyas myelin, acetylcholinesterase (AChE), non-phosphorylated neurofilament H (SMI-32), and the calcium-binding proteins parvalbumin (PV), calbindin (CB), and calretinin (CR) ^16, 20^. Additional information about laminar structure was assessed through the pattern of cross-frequency phase-amplitude coupling across SEF layers ^22^. Owing to variability in the depth estimates and the indistinct nature of the L6 border with white matter, some units appeared beyond the average gray-matter estimate; these were assigned to the nearest cellular layer.

### Acquiring eye position

Eye position data was collected at 1 kHz using an EyeLink 1000 infrared eye-tracking system (SR Research, Kanata, Ontario, Canada). This was streamed to a single data acquisition system (MAP, Plexon, Dallas, TX) and combined with other behavioral and neurophysiological data streams.

### Data collection protocol

The same protocol was used across monkeys and sessions. In each session, the monkey sat in an enclosed primate chair with their head restrained 45 cm from a CRT monitor (Dell P1130, background luminance of 0.10 cd/m^2^). The monitor had a refresh rate of 70 Hz, and the screen subtended 46° x 36° of the visual angle. After advancing the electrode array to the desired depth, they were left for 3 to 4 hours until recordings stabilized across contacts. This led to consistently stable recordings with single units typically held indefinitely. Once these recordings stabilized, an hour of resting-state activity in near-total darkness was recorded. This was followed by the passive presentation of visual flashes followed by periods of total darkness in alternating blocks. Finally, the monkey performed approximately 2000 trials of the saccade countermanding (stop-signal) task.

### Countermanding task

The countermanding (stop-signal) task utilized in this study has been widely used previously ^25^. Briefly, trials were initiated when monkeys fixated at a central point. Following a variable time period, drawn from an aging function to avoid anticipation of the visual stimulus ^40^, the center of the fixation point was removed leaving an outline. Simultaneously, a peripheral target was presented to the left or right of the screen.

On no stop-signal trials the monkey was required to shift gaze to the target. Fixation on the target was required for 600 ms, until an auditory tone sounded, whereupon monkeys could shift gaze anywhere. Fluid reward was delivered 600 ms later.

On stop-signal trials, comprising less than half of all trials, the center of the fixation point was re-illuminated after a variable stop-signal delay (SSD). An initial set of SSDs, separated by 40-60 ms for Monkey Eu and by 100 ms for monkey X, were selected for each recording session. To ensure that monkeys failed to countermand on ∼50% of stop-signal trials, SSD was adjusted through an adaptive staircasing procedure. When a monkey failed to inhibit a response, the SSD was decreased by 1, 2, or 3 steps (randomly drawn) to increase the likelihood of success on the next stop trial. When a monkey canceled the saccade, SSD was increased by 1, 2, or 3 steps (randomly drawn) to decrease the likelihood of success on the next stop trial. On stop-signal trials, the monkey was required to maintain fixation on the central point until the tone sounded, whereupon monkeys could shift gaze anywhere. Fluid reward was delivered 600 ms later. By design, the duration from target presentation until the tone was a fixed interval of 1500 ms. Thus, as SSD increased, the duration of fixation decreased (**Supplementary Figure 2b**).

Performance on this task is characterized by the probability of not canceling a saccade as a function of the SSD (the inhibition function) and the distribution of latencies of correct saccades in no-stop-signal trials and of noncanceled error saccades in stop-trials (**Fig 1b**). Performance of the stop-signal task is explained as the outcome of a race between a GO and a STOP process ^26^. The race model provides an estimate of the duration of the covert STOP process, the time taken to accomplish response inhibition, known as stop-signal reaction time (SSRT) ^29, 30, 79^. SSRT was calculated using two approaches—the conventional weighted-integration method and the more recent Bayesian Ex-Gaussian Estimation of Stop-Signal RT distributions (BEEST) ^86^ **(Supplementary Figure 3a, 4a, 5a).** Compared to weighted integration method, the Bayesian approach provides estimates of the variability in SSRT and the fraction of trigger failures for a given session ^86^. Individual parameters were estimated for each session. The priors were bounded uniform distributions (*μ_Go_*, *μ_Stop_: U* (0.001, 1000); *σ_Go_*, *σ_Stop_: U* (1, 500) *τ_Go,_ τ_Stop_: U* (1, 500); pTF: *U* (0,1)). The posterior distributions were estimated using Metropolis-within-Gibbs sampling ran multiple through three chains. We ran the model for 5000 samples with a thinning of 5. None of our conclusions depend on the choice of SSRT calculation method.

### Analysis of EEG

Methods paralleling those used in human studies were used. The N2 and P3 were obtained from average EEG synchronized on stop-signal presentation. Peak N2 was the time when the mean ERP reached maximal negativity in a 150-250 ms window after the stop-signal. Peak P3 was the time when the mean ERP in a 250-400 ms window after the stop-signal. The amplitude of the N2 and P3 was quantified as the mean Z-transformed voltage for each SSD in a ±50 ms window around the maximal ERP deflection determined for each session. Indistinguishable results were obtained with wider (±75 ms), and narrower (±25 ms) windows or just the instantaneous maximal polarization. To characterize the polarizations associated with response inhibition, a difference ERP (ΔERP) was obtained by subtracting from the ERP recorded on canceled trials the ERP recorded on RT-matched no stop-signal trials.

### Analysis of neural spiking

Spike density functions (SDF) for individual trials were constructed by convolving the spike times with a kernel matching the time course of an excitatory post-synaptic potential with an area equal to 1

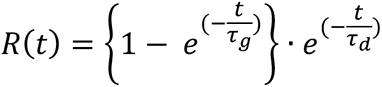

The influence of each spike (R(t)) increases with a short time constant (τ_g_ = 1 ms) and decays slower (τ_d =_ 20 ms). To analyze spiking activity associated with successful stopping, we compared the activity on canceled trials and on no stop-signal trials with RT greater than SSD + SSRT. This latency-matching compares trials in which countermanding was successful with trials in which countermanding would have been successful had the stop-signal been presented. Neurons were distinguished by patterns of modulation consisting of periods of facilitation or suppression using a consensus clustering algorithm ^28^ (**Supplementary Fig 1B**). The input to this analysis pipeline was the spike-density function on canceled trials and on latency-matched no stop-signal trials during the 100 ms preceding SSRT and 200 ms following SSRT. Results did not change much if interval durations were changed.

To prevent outlying values from exerting excessive influence, population spike density plots were obtained by scaling the SDF of each neuron by the 95% confidence interval between the 2.5% lowest rate and the 97.5% highest rate in one of two intervals. The first interval was a 600 ms window centered on SSRT on canceled and on no stop-signal trials. The second interval was −100 to +300 ms relative to the feedback tone.

To identify spiking modulation, we applied methods previously employed. First, we calculated a difference function (ΔSDF), the difference between the SDF on canceled and latency-matched no stop-signal trials. Periods of statistically significant modulation were identified based on multiple criteria—(a) the difference function must exceed by at least 2 standard deviations a baseline difference measured in the 100 ms interval before the target appeared, (b) the difference must occur from 50 ms before to 900 ms after the stop-signal, and (c) the difference must persist for at least 100 ms (or for 50 ms if the difference exceeded baseline by 3 standard deviations). As commonly found in medial frontal cortex, some neurons exhibited low spiking rates. To obtain reliable estimates of modulation times, we also convolved the SDF with a square 8 ms window. The modulation intervals were validated by manual inspection.

To determine modulation associated with the systematically variable timing of the feedback tone on canceled trials, the SDF was compared against the minimum value found between 500 ms before and 900 ms after the tone. Focusing on modulation occurring only during the period of operant control on behavior, modulations beginning less than 300 ms after the tone were not included. For comparisons across neurons and sessions, Z-transformed SDF or ΔSDF were used.

Spike widths of this sample of neurons exhibited a bimodal distribution ^16^. Consequently, neurons were distinguished as narrow- or broad-spikes. Narrow spike neurons had peak-to-trough duration less than 250 µs and broad spike, greater than or equal to 250 µs.

### Mixed effects models

We fit the variation in modulation of spiking or polarization of ERP to models of each of the behavioral and task measures as detailed in **Supplementary Figure 2**. We related neural modulation to the following models: (a) response conflict conceived computationally as the mathematical product of the activation of the race model GO and STOP processes and quantified as the probability of noncanceled error (P(error)) as a function of SSD, (b) P(error) contingent on viewing the stop-signal, denoted P(error | SS_seen_) and referred to as error likelihood, (c) absolute and log-transformed SSD, (d) hazard rate of stop-signal, (e) absolute and log-transformed delay until feedback tone, and (f) hazard rate of feedback tone. Although these behavioral and task measures can be correlated, random variations allowed for their differentiation.

To determine which performance measure accounted best for the variation of neural modulation, the performance and neural quantities were averaged within groups of early-, mid-, and late-SSD trials. SSD values greater than ∼350ms were not included because too few canceled trials were obtained. The analysis of the facilitation after SSRT as based on ΔSDF (**Fig 2, Fig 4**), but the major conclusions held if the analysis used SDF. The analysis of the modulation before SSRT or the feedback tone was based on the SDF of canceled trials. Before SSRT the SDF of canceled and no stop-signal trials was not different. Before the feedback tone, the interval was longer and more variable on canceled relative to no stop-signal trials.

Mixed-effects models of ΔSDF, SDF, or ΔERP values in relation to the various performance measures were compared using Bayesian Information Criteria (BIC). We report the results of the most basic version of each model with a main effect term corresponding to the performance parameter and random intercepts grouped by neuron (for spiking activity) or session (for ERP analysis). The values for each performance parameter were z-transform normalized for fair comparison between models related to different quantities. All constructed models had the same degrees of freedom, so BIC values between models could be compared directly. The model with the smallest BIC was endorsed as the best model. The fit of the other models relative to the best are reported using ΔBIC. As recommended ^87, 88^, ΔBIC (BIC_best_ – BIC_competing_) < 2 offers weak support against the competing model, 2 < ΔBIC < 6 offers strong support against the competing model, and ΔBIC > 6 conclusively rules out the competing model. More complex versions of these models resulted in similar conclusions. Mixed-effects models were performed using MATLAB’s Statistical Toolbox.

### Relating N2/P3 and neural spiking

We used the method described previously to establish the relationship between spiking activity and the ERN ^16^. Single trial spiking was the mean convolved spike data for that trial recorded from neurons in L2/3 and in L5/6 of perpendicular penetrations within ±50 ms of the N2 and P3 peaks. To account for variations in ERP voltage and spike counts across sessions, a fixed-effects adjustment was performed by centering each distribution on its mean and dividing by its most extreme value. To measure the N2/P3 amplitudes robustly, we grouped rank-ordered single-trial ERP values into 20 successive bins. From trials in each bin, we calculated the mean N2 and mean P3 magnitude (dependent variable), the mean spike count in the upper and lower layers (independent variables), and the average SSD, on Canceled trials. Data from all sessions were combined for a pooled partial correlation. Each point in **Fig. 5** plots the paired values of the mean normalized ERP voltage and normalized activity for each of the 20 bins from every session. The statistical relationship between ERP magnitude and spiking activity was quantified through multiple linear regression on normalized data pooled across sessions. Three factors were considered: (1) spiking activity in L2/3, (2) spiking activity in L5/6, plus (3) SSD to prevent its variation from confounding the relationship between ERP and neural spiking. However, as presented in the main text, the inclusion of this factor did not change the results.

## Code availability

The analysis codes that were used for this study are available from the corresponding author upon request.

## Data availability

The data that support the findings of this study are available from the corresponding author upon request.

## Acknowledgments

The authors thank G. Luppino, M. Matsumoto, N. Palomero-Gallagher, and, L. Rapan for sharing data; J. Elsey, M. Feurtado, M. Maddox, S. Motorny, J. Parker, D. Richardson, M. Schall, C.R. Subraveti, L. Toy, B. Williams, and R. Williams for animal care and other technical assistance; and Z. Fu, M. Matsumoto, P. Redgrave, U. Rutishauser, E. Sigworth, A. Tomarken, and G. Woodman for helpful discussions. Imaging data was collected in the Vanderbilt Institute of Imaging Science. This work was supported by R01-MH55806, R01-EY019882, P30-EY08126, Canadian Institutes of Health Research Post-Doctoral Fellowship, and by Robin and Richard Patton through the E. Bronson Ingram Chair in Neuroscience.

## Author contributions

Experimental design, J.D.S. Data collection, J.D.S. Data analysis, A.S. and S.E. Interpretation and preparation of manuscript, A.S., S.E., and J.D.S.

