## Supplementary Information for "Functional Architecture of Executive Control and Associated Event-Related Potentials"

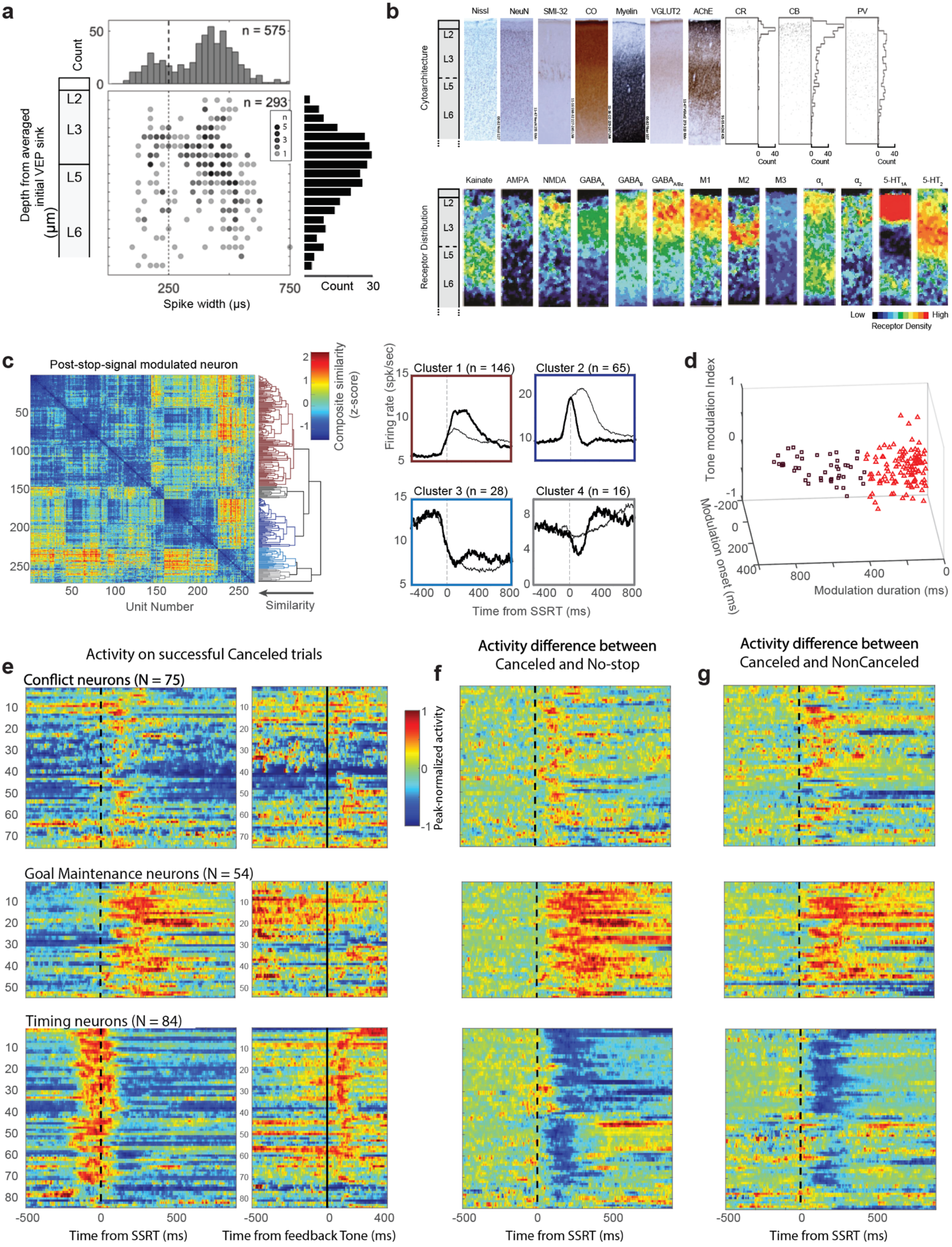
a
Count
n = 575
n = 293
Depth from averaged initial VEP sink (µm)
L2
L3
L5
L6
Spike width (µs)
Count
b
Nissl
NeuN
SMI-32
CO
Myelin
VGLUT2
AChE
CR
CB
PV
Cytoarchitecture
Receptor Distribution
Kainate
AMPA
NMDA
5-HT
Low
High
Receptor Density
c
Post-stop-signal modulated neuron
Composite similarity (z-score)
Unit Number
Similarity
Firing rate (spk/sec)
Cluster 1 (n = 146)
Cluster 2 (n = 65)
Cluster 3 (n = 28)
Cluster 4 (n = 16)
Time from SSRT (ms)
d
Tone modulation Index
Modulation onset (ms)
Modulation duration (ms)
Activity on successful Canceled trials
e
Conflict neurons (N = 75)
Goal Maintenance neurons (N = 54)
Timing neurons (N = 84)
Peak-normalized activity
Time from SSRT (ms)
Time from feedback Tone (ms)
Activity difference between Canceled and No-stop
f
Activity difference between Canceled and NonCanceled
g

**Supplementary Figure 1 |** SEF laminar structure and neuron classification.

**a,** Distributions of depths of units and spike widths sampled. Horizontal histogram shows the spike widths of all neurons sampled (n = 575), which exhibited bimodal values. The dashed line marks the 250 μs separation criterion used. Scatterplot shows variation of spike width across depth for neurons sampled in perpendicular penetrations (n = 293). The number of neurons at each time-depth indicated by gray scale (black = 5 neurons). The width of the spikes narrower than 250 μs does not vary with depth, and theincidence of encountering narrow spikes parallels the density of PV neurons [20]. The width of spikes wider than 250 μs increases from L3 to L6, which parallels the size of pyramidal neurons. Also, the incidence of isolated neurons decreases with depth, which parallels the density of pyramidal neurons. Vertical histogram shows the depths of all neurons sampled during the saccade countermanding task from the sessions with penetrations oriented perpendicular to the cortical layers. The color scale used for the time-depth plots in Figures 2, 3, and 4 indicates the number of neurons of each class sampled relative to the entire sampling distribution.

**b,** Laminar structure of SEF. Top row, from left to right are shown sections through SEF stained for Nissl, NeuN, nonphosphorylated neurofilament H (SMI-32), cytochrome oxidase (CO), Gallyas myelin, vesicular glutamate transporter 2 (VGLUT2), acetylcholinesterase (AChE), calretinin (CR), calbindin (CB), and parvalbumin (PV). Counts of CR, CB, and PV stained neurons are shown from[20]. In SEF, L1 is ~200 μm thick with some CR and CB but no PV neurons, and weak staining for CO, myelin, and VGLUT2. L2 is ~300 μm thick with dense stellate and few pyramidal neurons, the highest density of CR and CB neurons, but no SMI-32, stronger CO staining, fascicular myelin fibers, slightly stronger VGLUT2, and modest AChE. L3 is ~700 μm thick with a superficial sublayer with smaller pyramidal neurons and weak SMI-32 staining, very few CR, and modest densties of CB and PV neurons, stronger CO staining and denser myelin, VGLUT2, and AChE. A deeper sublayer is characterized by larger pyramidal neurons, pronounced SMI-32 staining, vanishingly few CR, less dense CB and modestly dense PV neurons, weaker CO, denser myelin and VGLUT2, and denser AChE. No granular L4 is evident in SEF. L5 is ~300 μm thick with large pyramidal neurons but inconsistent SMI-32 staining, no CR and fewer CB but modest density of PV neurons, lighter CO, denser myelin, lighter VGLUT2, and diminishing AChE staining. L6 is ~700 μm thick with smaller pyramidal neurons, light SMI-32, no CR, vanishingly few CB and low density of PV neurons, with lighter CO, still denser myelin, lighter VGLUT2 and sparse AChE staining. Lower row, sections processed for receptor autoradiography and color coded to visualize the laminar densities of various receptors from[23?]. The color scale maps to densities in fmol/mg protein with (minimum, maximum) levels as indicated for the following receptors: kainate (20, 1200), AMPA (20, 750), NMDA (200, 2500), $GABA_A$ (100, 3200), $GABA_B$ (150, 3500), $GABA_{A/Bz}$ (200, 3500), muscarinic $M_1$ (100, 1200), muscarinic $M_2$ (10, 350), muscarinic $M_3$ (100, 1200), adrenergic $\alpha_1$ (50, 700), adrenergic $\alpha_2$ (20, 800), serotinergic $5\text{-}HT_{1A}$ (20, 700), and serotinergic $5\text{-}HT_2$ (100, 500). Receptor densities reveal pronounced differences between L2/3 and L5/6 and other differences distinguishing L2 from L3 and L5 from L6. The variation in laminar structure can guide the investigation and interpretation of functional architecture of SEF.

**c,** Consensus clustering to distinguish patterns of modulation. In a sample of 575 neurons, we identified 213 with significant changes in spiking rate from presentation of the stop-signal until 200 ms after stop-signal reaction time (SSRT) on successfully canceled trials compared to latency-matched no-stop trials. The spike density function on canceled and no-stop trials in intervals 0–100 ms before and 0–200 ms after SSRT were submitted to an unsupervised consensus clustering pipeline [27]. The matrix and associated dendrogram in left panel illustrate, with color coding, the outcome of the clustering algorithm. Note that the colors used in this figure bear no relation to those used elsewhere in the manuscript. The heatmap indicates the z-score of a composite similarity measure. Of 5 resulting clusters, only 4 included enough neurons to analyze further. Among these 4, neurons clustered along two dimensions. First, neurons were distinguished by a relative increase (left spike-density plots) or decrease (right spike-density plots) in discharge rate on canceled (thick spike density function) compared to latency-matched no-stop trials (thin spike density function). Second, neurons were distinguished by facilitation (top) or suppression (bottom) around SSRT. Cluster 1 (upper left) included 146 neurons; cluster 2 (upper right), 65 neurons; cluster 3 (lower left), 28 neurons; and cluster 4 (lower right), 16 neurons. Neurons in clusters 1 and 2 were analyzed further. The clustering results were verified by manual curation, resulting in some cluster 3 and 4 neurons being included in clusters 1 and 2 and removal of neurons with poor signal quality.

**d,** Neurons with facilitated activity were distinguished further using k-means clustering. In the 3D scatter plot, each neuron is represented by a point marking modulation duration, modulation latency, and tone modulation. Tone modulation was the contrast (difference / sum) between spike counts 200 ms before and 200 ms after the feedback tone. The values along the three dimensions were normalized for K-means clustering (the figure shows the true values on the axes). We considered 2, 3, 4, and 5 clusters but used just 2 based on qualitative examination of population spike density functions. The facilitated neurons were mainly differentiated by modulation duration.

**e,** Heat-maps representing normalized spike-density function on successful canceled trials for all neurons reported in this paper. The spike-density function for each neuron is normalized to its peak activity ±300 ms around SSRT. The left panel plots activity aligned on stop-signal reaction time (SSRT) (dashed black line). The right panel plots activity aligned on the feedback tone (solid line). Each row corresponds to activity of one neuron with higher discharge rates in hotter, and lower discharge rates in cooler colors. The time course of activation that distinguished conflict, event-timing, and goal maintenance neurons is evident in the three pairs of panels. Similarities and differences between these SEF neurons and DA neurons in SNpc and neurons in striatum sampled during saccade countermanding is afforded by comparison with Figures 2 and 5 of Ogasawara et al. (2018) [45].

**f,** Heat-maps representing the difference in spike-density function on successful canceled trials relative to latency-matched no-stop trials, aligned on SSRT, selected for the early stop-signal delay (SSD). The difference function for each neuron is normalized to the maximum deviation. Hot colors represent facilitation and cold colors represent suppression on canceled relative to no-stop trials. The patterns of activation that distinguish the three major categories of neurons are more evident across the three panels. Similarities and differences between these SEF neurons and DA neurons in SNpc and neurons in striatum sampled during saccade countermanding is afforded by comparison with Figures 2 and 5 of Ogasawara et al. (2018) [45].

**g,** Heat-maps representing the difference in spike-density function on successful canceled trials relative to non-canceled trials (in spite of the inherent difference in underlying RT), aligned on SSRT, selected for an intermediate SSD with enough trials of both types. The patterns of activation that distinguish the three major categories of neurons are more evident across the three panels. Similarities and differences between these SEF neurons and DA neurons in SNpc and neurons in striatum sampled during saccade countermanding is afforded by comparison with Figures 2 and 5 of Ogasawara et al. (2018) [45].

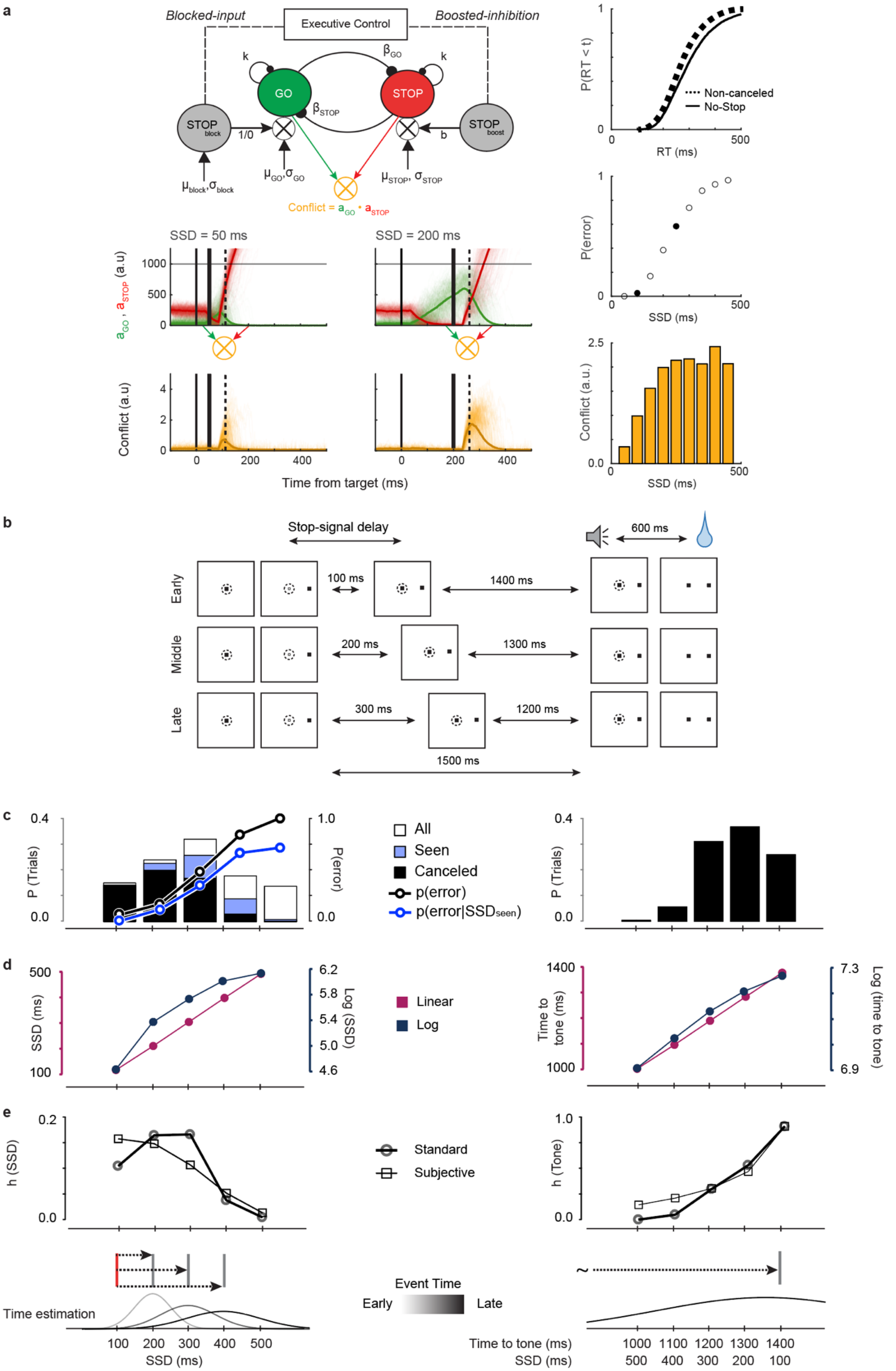
a
Blocked-input
Executive Control
Boosted-inhibition
k
βGO
k
GO
STOP
βSTOP
STOP block
1/0
b
STOP boost
μblock, σblock
μGO, σGO
μSTOP, σSTOP
Conflict = aGO • aSTOP
SSD = 50 ms
SSD = 200 ms
aGO , aSTOP (a.u)
1000
500
0
Conflict (a.u)
4
2
0
0
200
400
Time from target (ms)
P(RT < t)
1
0
0
500
Non-canceled
No-Stop
RT (ms)
P(error)
SSD (ms)
Conflict (a.u.)
2.5
0.0
SSD (ms)
b
Stop-signal delay
600 ms
Early
100 ms
1400 ms
Middle
200 ms
1300 ms
Late
300 ms
1200 ms
1500 ms
c
0.4
0.0
P (Trials)
1.0
0.0
P(error)
All
Seen
Canceled
p(error)
p(error|SSDseen)
d
500
100
SSD (ms)
6.2
5.8
5.4
5.0
4.6
Log (SSD)
Linear
Log
1400
1000
Time to tone (ms)
7.3
6.9
Log (time to tone)
e
0.2
0.0
h (SSD)
Standard
Subjective
1.0
0.0
h (Tone)
Time estimation
Event Time
Early
Late
100 200 300 400 500
SSD (ms)
Time to tone (ms)
SSD (ms)
1000 1100 1200 1300 1400
500 400 300 200 100

**Supplementary Figure 2 |** Parameters tested for neural modulation.

The analysis and interpretation of patterns of neural spiking were based on multiple computational measures. We considered response conflict and the associated error likelihood. We also considered the temporal statistics of the key task events: presentation of the stop-signal after stop-signal delay (SSD), and presentation of the feedback tone after a variable interval after SSRT on canceled trials. Below, we describe the set of observed and inferred task parameters tested to determine the coding scheme of neural signals reported in this study.

**a, Conflict during response inhibition**. Its formal computational definition as simply the product of the co-activation of competing response channels or processes was a chief virtue of the original formulation of response conflict [30]. That definition translates naturally into saccade countermanding through the interactive race model [28, 29]. Here we demonstrate that translation.

**Left:** Interactive race architectures. Excitatory connections terminate with arrowheads, and inhibitory connections terminate with circles. Response production or inhibition is governed by the stochastically evolving state of a GO unit and a STOP unit. A response is produced if the GO unit activation reaches a threshold. The response is canceled if inhibition from the STOP unit prevents it from reaching threshold. Stochastic activation of the GO unit begins an afferent delay after the target appears and is driven by an excitatory input (average $\mu_{GO}$ ± standard deviation $\sigma_{GO}$) with leak ($k_{GO}$). The STOP unit is driven by an excitatory input ($\mu_{STOP}$ ± $\sigma_{STOP}$) with leak ($k_{STOP}$) from before target presentation. After target presentation, inhibition from the GO unit ($\beta_{GO}$) reduces STOP unit activation. Two mechanisms have been proposed whereby the STOP unit can interrupt GO unit activation. Both instantiate delayed potent interruption, allowing the network of interacting units to produce behavior that can be described as the outcome of a race between stochastically independent processes [25]. First, through boosted-inhibition, the STOP unit becomes active an encoding delay after the stop-signal is presented, and the inhibition ($\beta_{STOP}$) from the STOP unit is amplified by a multiplicative boost (b) potently reduces activation of the GO unit. Second, through blocked-input, activation of the STOP unit blocks the input to the GO unit, whereupon activation is reduced through leak.

Both models make explicit the need for an executive control process that enables either the boost or the block (or both) to re-balance potently the state of the network. The need for such executive control indicates that the un-balanced state of the network reacting to the stop-signal is unusual, as the unexpected interruption of a motor plan typically is. The GO and STOP units are competing response channels, so the product of the co-activation of the GO and STOP units is a measure of response conflict in this task. This was quantified in simulations with parameters sufficient to simulate observed RT and P(Error) values (right panels). Individual (thin) and average (thick) activations of the GO (green) and STOP (red) units on simulated canceled trials with SSD = 50 ms (left) and SSD = 200 ms (right). The activation of the GO unit increases closer to the threshold (gray) on progressively longer SSD. Consequently, interruption of GO unit activation by the STOP unit fails more often on progressively longer SSD. Individual (thin) and average (thick) plots of the instantaneous product of $a_{GO}$*$a_{STOP}$ are plotted below. The conflict product arises at and peaks after SSRT.

**Right:** Top, simulated RT on trials with no stop-signal (thin solid) and non-canceled errors (thick dotted). Middle, simulated inhibition function with the two illustrated SSD distinguished. Bottom, average conflict as a function of SSD from simulated trials. The results of this simulation demonstrate that the product of the activation of GO and STOP units that produce countermanding performance provides directly a measure of conflict that is proportional to P(Error). Consequently, P(Error) is an effective proxy for the conflict measure.

**b, Timing of trial events**. Using conventions in Figure 1, trials with 100, 200, and 300 ms SSD are portrayed. By design, on canceled trials, the interval between presentation of the visual target and of the feedback tone was fixed at 1,500 ms with another fixed 600 ms between the tone and fluid reward delivery. Consequently, on canceled trials, the interval from stop-signal to success tone ($T_{tone}$) decreased linearly with SSD because $T_{tone}$ = 1500 – SSD. Based on previous findings that SEF neurons signal event timing and interval duration[13,14] we determined whether the neurons we sampled signal the temporal statistics of SSD or the delay to the feedback tone that signals when operant control ends and reward will be delivered.

**c,** Monkeys experienced stop-signals at a range of times. Saccades were canceled for shorter SSD. At longer SSD the monkey may not see the stop-signal. After SSRT on canceled trials, monkeys next experienced a delay until a feedback tone, which co-varied with SSD. To ensure that monkeys failed to cancel on ~50% of all stop-signal trials, SSD was adapted dynamically during data acquisition in a staircase algorithm. After non-canceled error trials, SSD was decreased. After canceled trials, SSD was increased. This adjustment happened even if the stop-signal was not seen and resulted in fewer trials with shorter or longer SSD. To discourage monkeys from exploiting the temporal statistics of SSD[32], SSD was adjusted in steps of 1, 2, or 3 intervals randomly selected with uniform 1/3 probability. This is illustrated in lower left panel. After a canceled trial with SSD of 100 ms (red), the SSD on the next trial would increase to either 200, 300, or 400 ms. **Left panel.** With left ordinate, the distribution of stop-signal trials in which the saccade was successfully canceled (black), those in which the stop-signal was seen ($SS_{seen}$; blue) and all stop-signal trials (open). With right ordinate, the P(Error) (black) and P(error | $SS_{seen}$) (blue) as a function of SSD. Inspired by reinforcement learning models, we determined whether neural spiking following the stop-signal represents just P(Error) or P(error | $SS_{seen}$), which is a proxy of error likelihood learned through experience with the task. Thus, while correlated in this task, conflict (P(Error)) and error likelihood (P(error | $SS_{seen}$)) diverge at longer SSDs in which many non-canceled errors are generated before stop-signal appears (i.e., RT < SSD). Inspired further by reinforcement learning models and the necessity of enforcing the canceled saccade state for an extended period, we also determined whether neural spiking encoded the temporal statistics inherent to this task. **Right panel,** Distribution of intervals between SSRT and the feedback tone. Knowledge of the time of the feedback tone relative to stop-signal can be acquired only on canceled trials. For no-stop and non-canceled error trials, the interval until the feedback tone had no variability.

**d,** Based on previous research[34], we determined whether neural signals represented absolute or a log-compressed intervals. **Left panel.** With left ordinate, SSD, and right ordinate, log (SSD). **Right panel.** With left ordinate, $T_{tone}$, and right ordinate, log($T_{tone}$).

**e,** The instantaneous probability or expectation of the occurrence of an event is quantified through the hazard rate: h(t) = f(t)/[1-F(t)], where f(t) is the probability density, and F(t) is the cumulative distribution of event times. The hazard rate model would test if neural signals preceding an event represents the instantaneous probability (or expectation) of the event [34] or if neural signals following an event relate to the surprise upon the violation of that expectation [12, 37].

**Left panel.** Hazard rate (circle) and expectation (square) of SSD. The calculation of these quantities entailed two kinds of nuance. First, because stop-signal appearance is probabilistic (~40-50% likelihood), the hazard function is calculated with the conditional probability of stop-signal appearance for each session. This conditional probability accounted for the fact that on a given trial, because of the uncertainty associated with the current trial being a stop-signal trial, if time passes and no stop-signal is observed, the belief state about the current trial being a stop-signal trial drops, reducing the expectation for stop-signal appearance. Because our approach relied on mean responses across many trials having different patterns of preceding trial history within each SSD bin, we assumed variations in belief state due to passage of time only within a trial. We also considered the possibility that knowledge of the temporal structure of SSD can be acquired from all stop-signal trials in which stop-signal was seen (blue and black bars), or solely from canceled trials (black bars). Second, because subjective estimation of intervals is not accurate or precise, instantaneous expectation can diverge from actual instantaneous probability. The estimation of elapsed time has increasing uncertainty over longer intervals[65]. To account for inaccuracy in time estimation, we convolved f(t) with a Gaussian kernel with standard distribution proportional to the elapsed time:

$$g(t) = \frac{1}{\emptyset t\sqrt{2\pi}} \int_{-\infty}^{\infty} f(t) e^{-\left(\frac{\tau - t}{\sqrt{2}\emptyset t}\right)^2} dx$$

The coefficient of variation, ø, was set to 0.26 based on previous research [34, 65]. We also calculated trial-wise hazard rate with f(t) changing according to trial history (not shown).

**Right panel.** Hazard rate (circle) and expectation (square) of $T_{tone}$. Unlike the stop-signal, the tone was deterministic (100% likelihood) with temporal statistics learned only on canceled trials. We considered the standard hazard rate based on the overall distribution of feedback time, the subjective hazard rate, accounting for the increased inaccuracy in subjective estimation as a function of elapsed time, and the dynamic hazard rate, that accounted for the knowledge of the underlying relationship between SSD and feedback time. Because of the linear relationship between SSD and feedback time on canceled trials, for the calculation of the dynamic hazard rate (not shown), the experienced SSD restricted the feedback time to only a single value. This is schematically depicted in the lower panel (right) for the same experienced SSD on the left panel. this value was then smoothed with the Gaussian kernel as described above before hazard rate was calculated. The tone became increasingly likely to happen with elapsed time as an aging distribution.

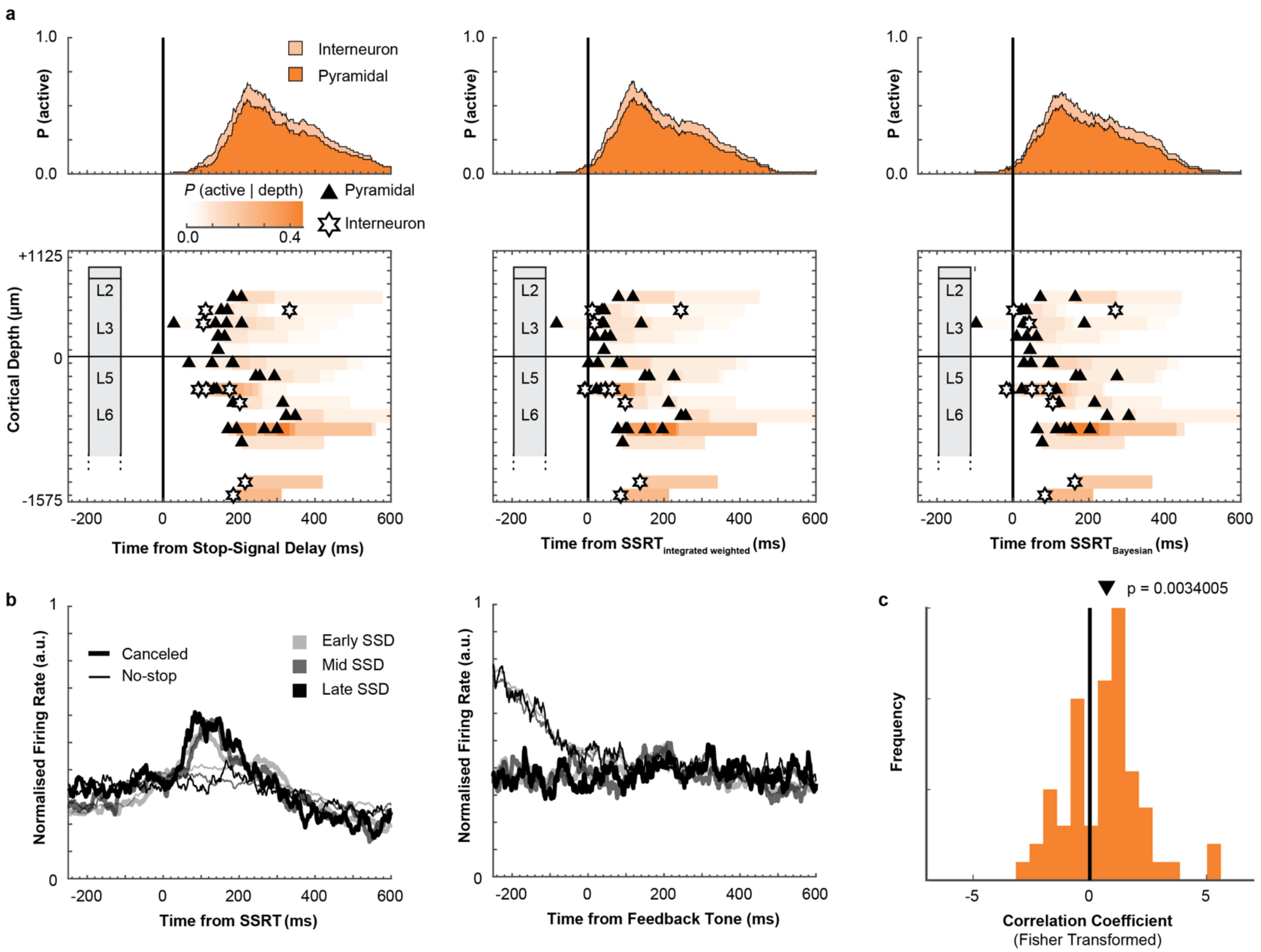


**Supplementary Figure 3 |** Conflict Neurons.

**a,** Recruitment and time-depth plots aligned on presentation of the stop-signal (left), SSRT calculated based on the method of integration (middle, same as **Fig 2**), and the Bayesian Estimation of Ex-Gaussian Stop-Signal reaction time (BEESTS) (right). The patterns supporting the conclusions reported in the main text do not vary with the method of alignment. This neural signal predominantly follows SSRT.

**b,** Normalized population spike-density functions aligned on SSRT (left) and feedback time (right) for canceled (thick) and RT-matched no stop trials (thin) subsampled from trials with low (lightest), middle (intermediate), and high (darkest) P(Error). The scaling of the modulation with SSD and P(Error) is evident across the sample.

**c,** Distribution of correlation coefficients of post-SSRT modulation as a function of P(Error) across sampled neurons. Correlation coefficients were transformed into a normal distribution using Fisher-transformation. The distribution of correlation coefficients were shifted significantly in the positive direction (t-test, t (63) = 3.04, p = 0.0034). The black triangle indicates the mean of the distribution. Viewed in this format, the correspondence is clear between the modulation pattern of SEF and that of dopamine neurons in SNpc of monkeys performing a saccade countermanding task [45].

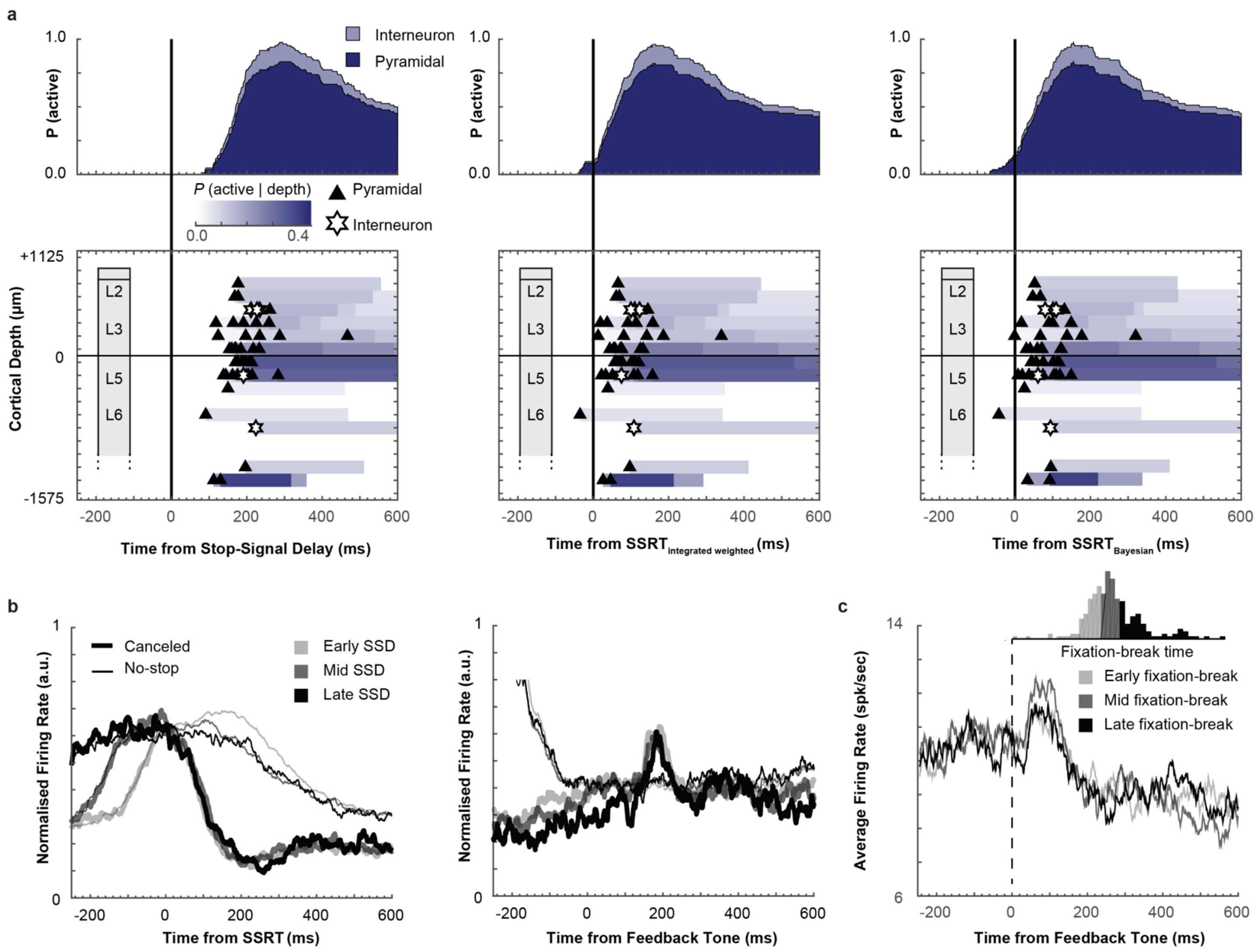


**Supplementary Figure 4 |** Event-Timing Neurons

**a,** Recruitment and time-depth plot aligned on stop-signal (left), SSRT calculated based on the method of integration (middle, same as **Fig 3**), and the Bayesian Estimation of Ex-Gaussian Stop-Signal reaction time (BEESTS) (right). The patterns supporting the conclusions reported in the main text do not vary with the method of alignment. This neural signal predominantly begins after SSRT and persists until the tone.

**b,** Normalized population spike-density functions aligned on SSRT (left) and feedback time (right) for canceled (thick) and RT-matched no stop trials (thin) subsampled from trials with low (lightest), middle (intermediate), and high (darkest) P(Error). The timing of the modulation across SSD is evident across the sample.

**C,** Population spike-density function (n = 84) aligned on feedback tone for short (light gray), medium (dark gray), and long (black) periods until fixation broke from a central point after the tone. Inset figure plots distribution of the time of the first saccade or blink following the feedback tone in one session. The temporal dynamics of the tone-aligned activity did not depend on the time at which fixation was broken following the feedback tone. This pattern was observed on almost all individual neurons (not shown).

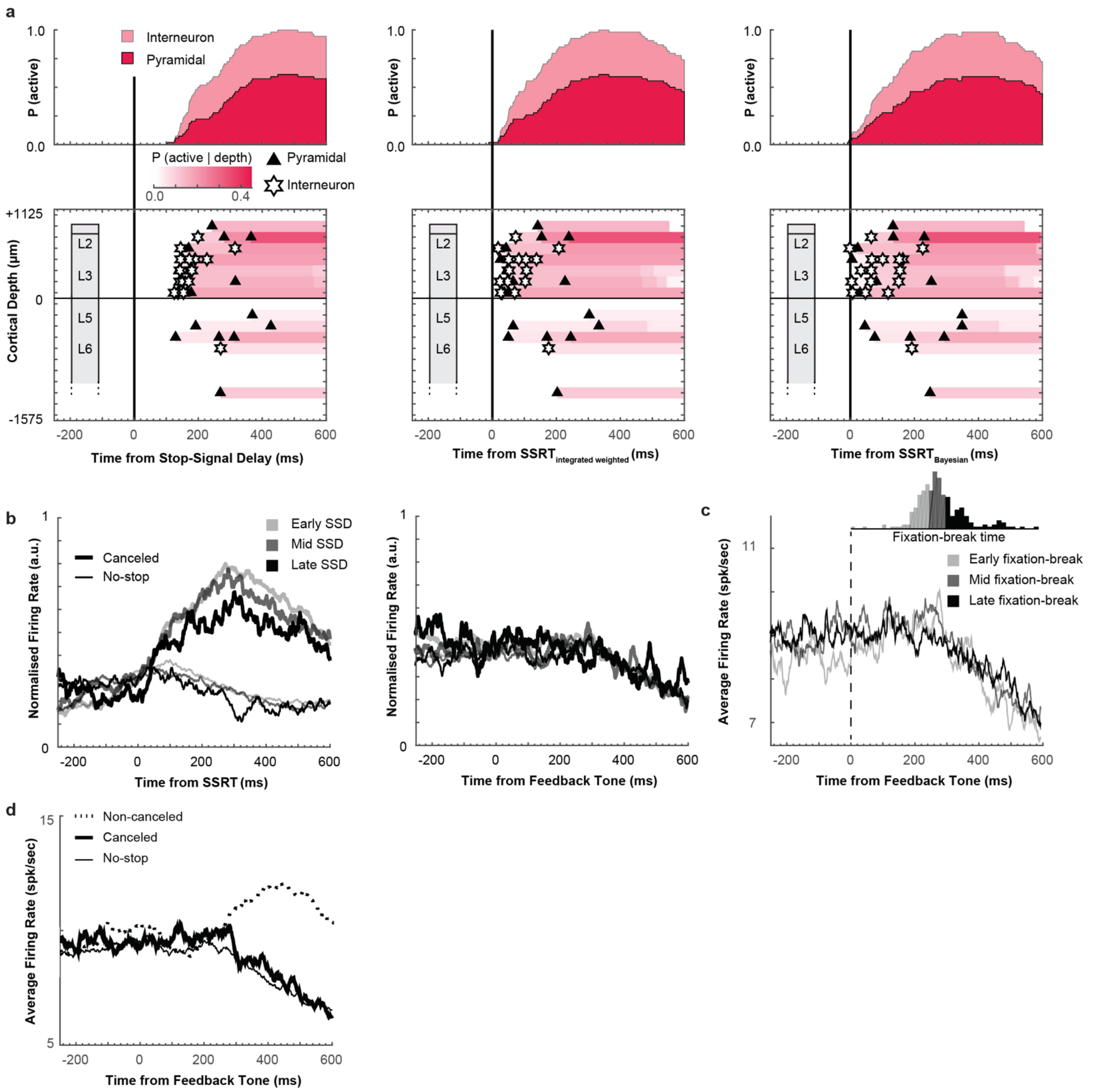


**Supplementary Figure 5 |** Goal Maintenance Neurons

**a,** Recruitment and time-depth plots aligned on stop-signal (left), SSRT calculated based on the method of integration (middle, same as **Fig 4**), and the Bayesian Estimation of Ex-Gaussian Stop-Signal reaction time (BEESTS) (right). The patterns supporting the conclusions reported in the main text do not vary with the method of alignment. This neural signal grows after SSRT and persists until after the tone.

**b,** Normalized population spike-density functions aligned on SSRT (left) and feedback time (right) for canceled (thick) and RT-matched no stop trials (thin) subsampled from trials with low (lightest), middle (intermediate), and high (darkest) P(Error). The scaling of the modulation across SSD is evident across the sample.

**c,** Population spike-density function (n = 54) aligned on feedback tone for short (light gray), medium (dark gray), and long (black) periods until monkeys shifted gaze from the central point or blinked (inset plots distribution of the time of the first saccade or blink following the feedback tone in one session). The temporal dynamics of the tone-aligned activity did not depend on the time at which fixation was interrupted following the feedback tone. This pattern was observed on almost all individual neurons (not shown).

**d,** Population spiked-density function on canceled (thick solid), no-stop (thin), and non-canceled (thick dashed) trials. A large proportion of Goal Maintenance neurons were also classified as Loss neurons, with higher activity on unrewarded compared to rewarded trials.

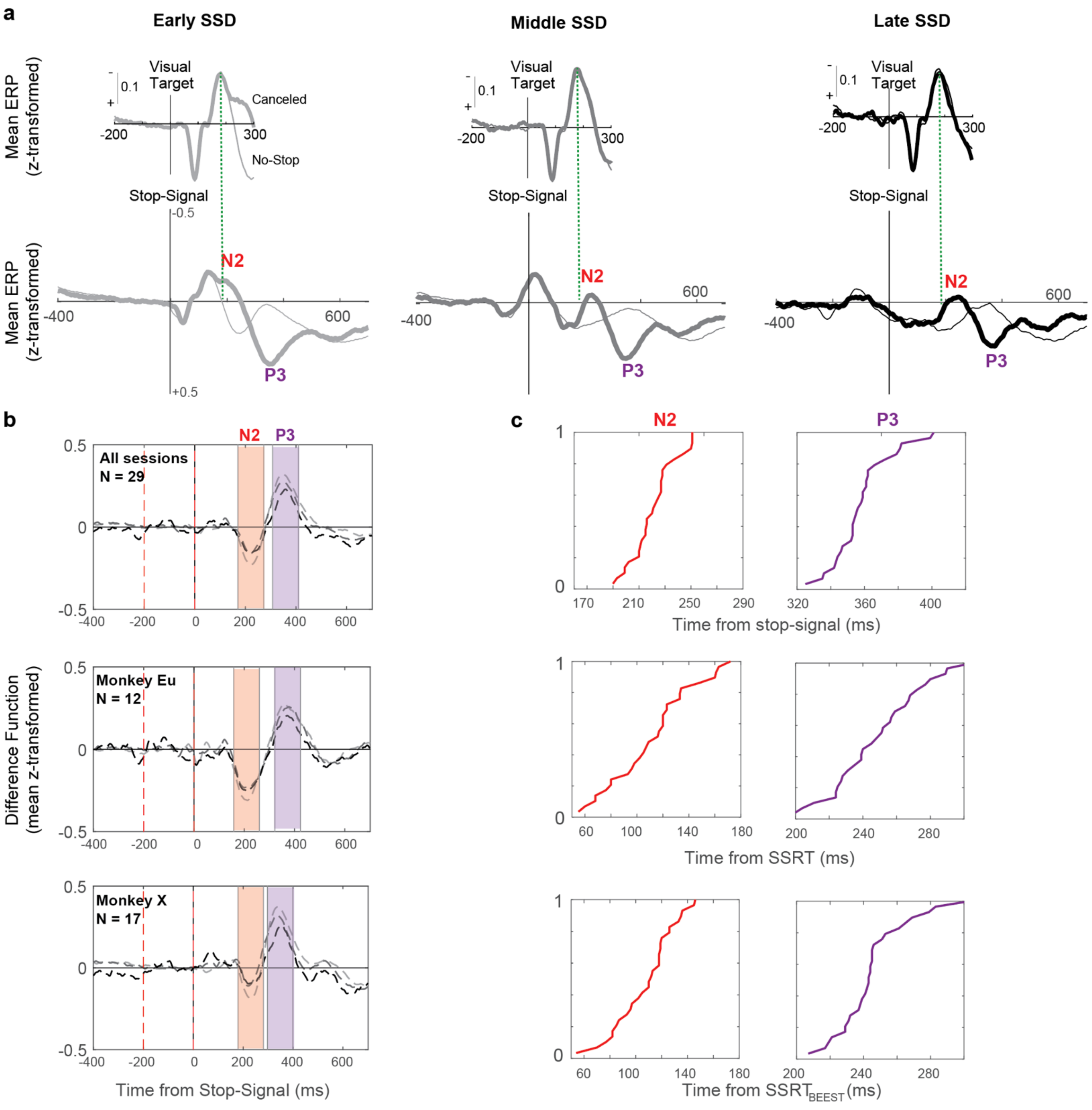


**Supplementary Figure 6 |** N2/P3 Characteristics

**a,** ERP polarization aligned on visual target presentation (top), and stop-signal (bottom) for early (left), intermediate (middle), and late (right) SSD associated with lower, intermediate, and high P(error). At all SSD, the visually-evoked negativity (green dashed line) occurs earlier than the peak of N2 by ~ 40 ms. This is evidence that the N2 is not just a visual response.

**b,** N2 and P3 shown for all sessions (top), sessions from monkey Eu (middle) and X (bottom). The N2 and P3 exhibited similar features across monkeys. The shaded intervals mark a ±50 ms period around the peak of each ERP.

**c,** Cumulative distributions of N2 (left) and P3 (right) peak times aligned on stop-signal (top), SSRT by method of integration (middle) and SSRT by BEESTS (bottom). The distributions are narrower when aligned on stop-signal, suggesting better time-locking of the peak of these ERPs to stop-signal.

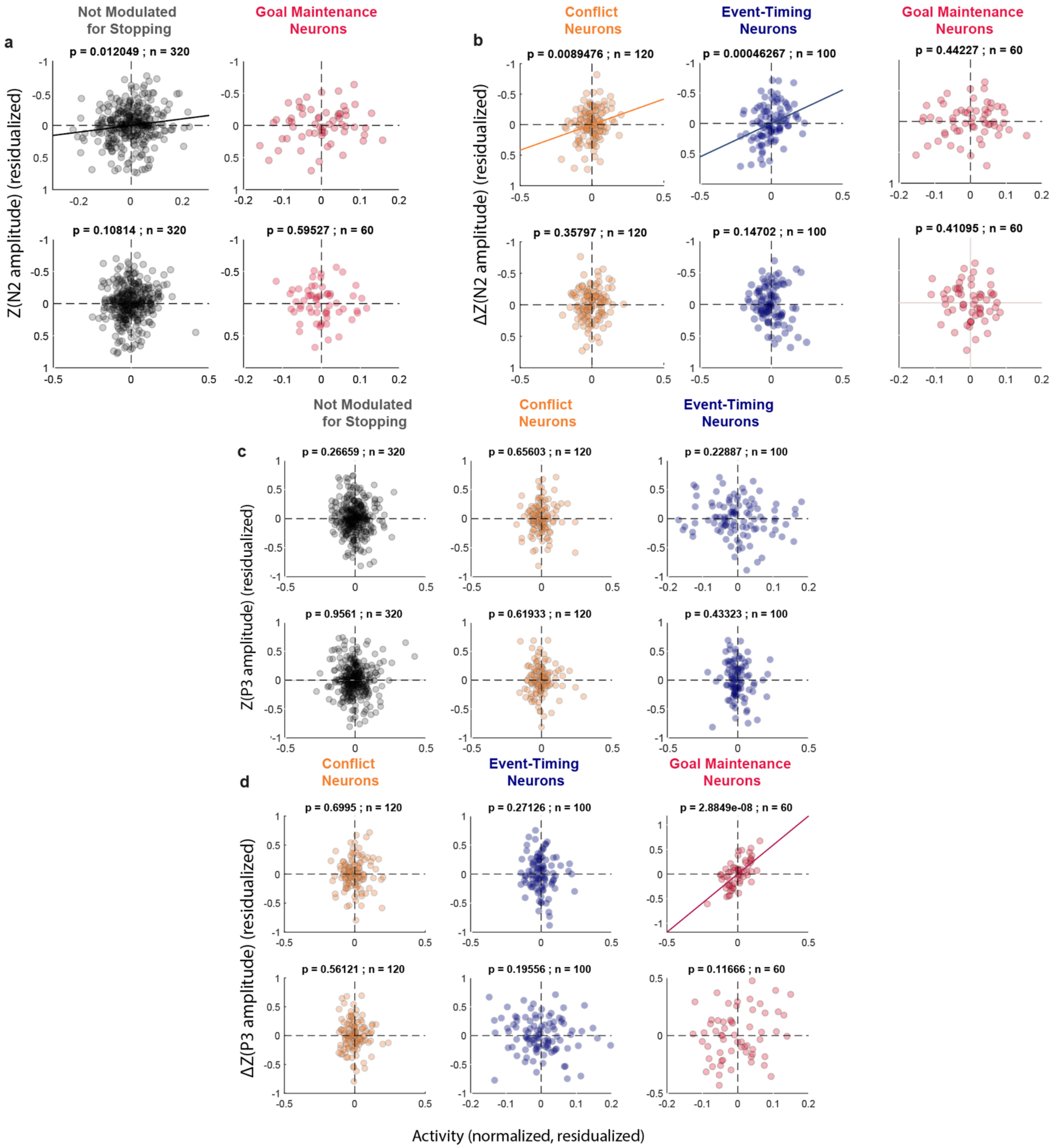


**Supplementary Figure 7 |** N2/P3 relationship with spiking

**a,** N2 relationship with neuron classes not shown in **Fig 5e** and **Fig 5f**. N2 polarization was predicted by the spiking of other SEF neurons that were not modulated around the time of SSRT in L2/3 but not L5/6. N2 polarization was unrelated to the spiking of Goal Maintenance neurons in L2/3 or L5/6.

**b,** Results of correlation analysis of N2 polarization as a function of spiking activity of the different types of neurons performed by subtracting polarization on latency-matched no-stop trials. The results are effectively identical to those obtained from measurement of simple N2 polarization.

**c and d**, These plots show the parallel analyses of the period of P3 polarization shown in panels **a** and **b**. The P3 is associated only with the spiking of Goal Maintenance neurons in L2/3.

6

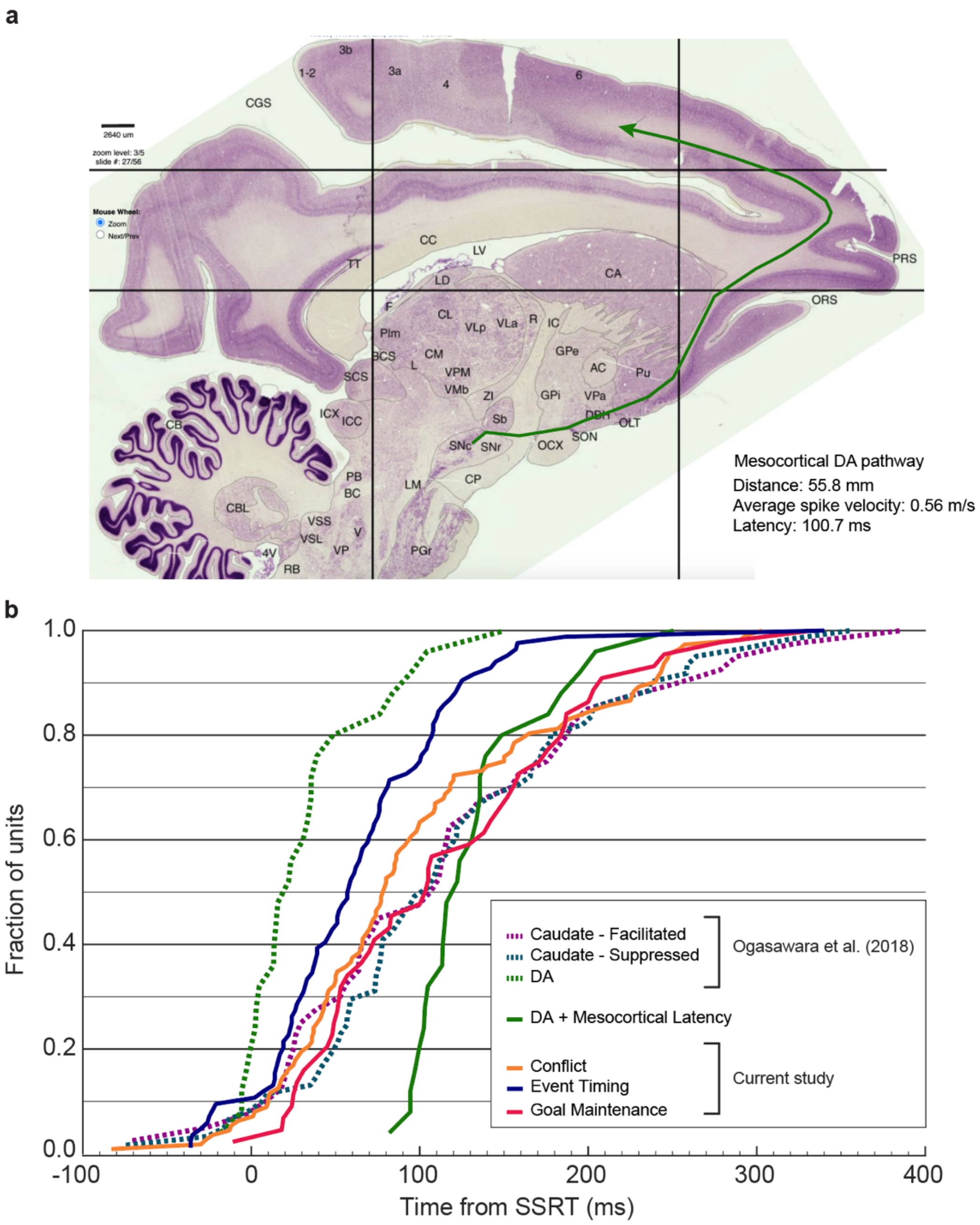


c

| | Conflict (97.8 ± 7.8 ms, n = 112) | Event Timing (62.4 ± 6.4 ms, n = 84) | Goal Maintenance (114.8 ± 11.9 ms, n = 44) |
|---|---|---|---|
| **SNpc** (32.6 ± 8.1 ms, n = 25) | p = 0.002 * | p = 0.602 | p < 0.001 * |
| **SNpc + mesocortical latency** (128.7 ± 8.1 ms, n = 25) | p = 0.531 | p = 0.003 * | p = 0.991 |
| **Caudate (Facilitation)** (114.4 ± 15.8 ms, n = 40) | p > 0.999 | p = 0.008 * | p > 0.999 |
| **Caudate (Suppression)** (116.2 ± 11.3 ms, n = 61) | p = 0.737 | p = 0.001 * | p > 0.999 |

One-way ANOVA, Neuron Type x Onset Latency, $F(6, 384) = 7.87$, $p < 0.001$
Tukey post-hoc comparisons (* = significant at the $p < 0.05$ level)

**Supplementary Figure 8 |** Relationship of SEF modulation to substantia nigra pars compacta and caudate nucleus.

**a,** The mesocortical pathway of macaque monkeys. To infer the temporal relationship between VTA/SNpc and SEF signals, the mesocortical pathway was traced on a sagittal slice from the RH12 dataset (Dataset 6, RH12, Slide 27/56, Sagittal, Nissl; brainmaps.org). The estimated was 55.8 mm. Based on measurements of the conduction velocity of DA axons in rodents and assuming similar values for primates (0.54 m/s[47]; 0.55 m/s[48]; 0.58 m/s[49]), we estimate that the conduction time of a spike from SNpc to SEF is 100.7 ms (average of 103.4 ms, 101.5 ms, 96.3 ms).

**b,** Modulation times of the SEF neurons and of dopaminergic and striatal neurons. Cumulative distributions conflict (gold), event timing (blue), and goal maintenance (red) neurons in SEF are plotted with the latencies of dorsolateral SNpc neurons (dashed green) and of facilitated (dashed purple) and suppressed (dashed blue) neurons in the head of the caudate nucleus obtained from [45]. Also plotted is the estimated distribution of DA spike arrival times in SEF based on the conduction time from SNpc to SEF (green).

**c,** Comparisons of modulation times. Mean ± SEM latencies for type of neurons are shown beneath. A one-way independent measures ANOVA showed significant differences in modulation latencies between the different neuron types in SEF and the SNpc and caudate neurons ($F(6,384) = 7.87$, $p < 0.001$). The table reports outcomes of Tukey post-hoc comparisons with statistically significant differences marked by an asterisk. SNpc DA neurons modulated significantly earlier than conflict ($p = 0.002$) and goal maintenance ($p < 0.001$) neurons but were not different from event timing neurons ($p = 0.602$). However, the estimated arrival times of DA spikes in SEF were not different from the modulation times of conflict ($p = 0.531$) or goal maintenance ($p = 0.991$) neurons but were significantly later than the modulation of event timing neurons ($p = 0.003$). Meanwhile, both facilitation and suppression after SSRT in the caudate nucleus arose significantly later than the modulation of event timing neurons in SEF (facilitation $p = 0.008$; suppression $p < 0.001$) but simultaneously with conflict (facilitation $p > 0.999$; suppression $p = 0.737$) and goal maintenance (facilitation $p > 0.999$; suppression $p > 0.999$) neurons.

| Site | n. penetration | All neurons | Conflict | Event Timing | Goal Maintenance |
|---|---|---|---|---|---|
| np1 | 6 | 140 | 10 | 29 | 12 |
| np2 | 7 | 142 | 29 | 6 | 8 |
| P1 | 6 | 104 | 12 | 41 | 1 |
| P2 | 6 | 133 | 18 | 6 | 30 |
| P3 | 4 | 56 | 6 | 2 | 3 |
| Total count | | 575 | 75 | 84 | 54 |
| $X^2(4, 575)$ | test statistics | - | 11.62 | 84.13 | 39.3 |
| | p-value | - | 0.0204 | < $10^{-5}$ | < $10^{-5}$ |

| Layer | | All neurons | Conflict | Event Timing | Goal Maintenance |
|---|---|---|---|---|---|
| L2 | | 34 | 2 | 3 | 7 |
| L3 | | 114 | 11 | 22 | 19 |
| L5 | | 88 | 14 | 19 | 6 |
| L6 | | 39 | 7 | 2 | 1 |
| L6+ | | 18 | 2 | 3 | 1 |
| Total count | | 293 | 36 | 49 | 34 |
| $X^2(4, 293)$ | test statistics | - | 4.28 | 11.24 | 7.33 |
| | p-value | - | 0.369 | 0.024 | 0.12 |

**Supplementary Table 1 | Distribution of Conflict, Event Timing, and Goal Maintenance signals across recording sites and cortical depths**

**Top,** Count of Conflict, Event Timing, and Goal Maintenance signals across the 575 recorded neurons sampled from five sites in monkey Eu and monkey X. Three of the five sites were sampled with perpendicular penetrations (monkey Eu: P1, monkey X: P2 and P3) and two were not (monkey Eu: np1, monkey X: np2). The bottom row shows the test statistics for homogeneity based on a chi-square test. For each signal a 5-by-2 contingency matrix was constructed based on the counts of units recorded at each site (5 rows) with or without a given response type. The probability of sampling each kind of signal varied significantly across sites.

**Bottom,** Count of the neural signals from sites P1, P2 and P3 across cortical depth. Contacts ranged from depth 1 (surface) to depth 19 (deepest). Depths 2-16 span the estimated cortical gray matter. The alignment of sampling depths across sessions has been previously described [16, 20]. Data is divided into the four major cortical layers plus contacts deemed to lie beyond the L6 boundary. The bottom row shows the test statistics for homogeneity based on a chi-square test. While Conflict neurons and Time Keeping neurons were distributed equally across cortex, Goal Maintenance neurons were found significantly more often in L2/3 relative to L5/6.

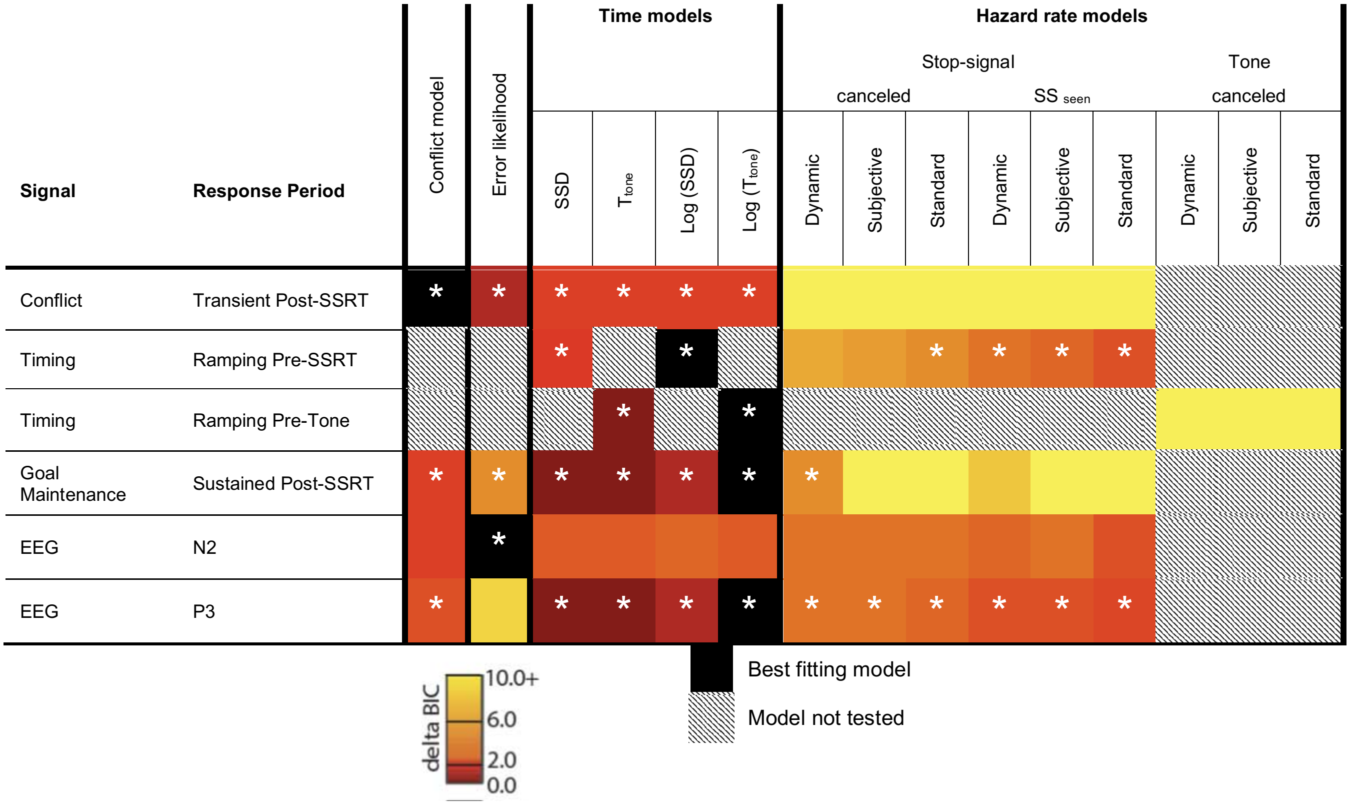


| Signal | Response Period | Predictor | df | β | SE | *t-statistics* | p |
|---|---|---|---|---|---|---|---|
| Conflict | Transient Post-SSRT | Intercept | 104 | 2.66 | 0.31 | 8.47 | 1.68E-13 |
| | | P(Error) | | 0.78 | 0.22 | 3.57 | 0.000535 |
| Goal Maintenance | Sustained Post-SSRT | Intercept | 152 | 6.68 | 0.73 | 9.09 | 5.26E-16 |
| | | Log ($T_{tone}$) | | 0.52 | 0.15 | 3.53 | 0.000557 |
| Timing | Ramping Pre-SSRT | Intercept | 250 | 16.19 | 1.28 | 12.62 | 1.41E-28 |
| | | Log (SSD) | | 0.76 | 0.24 | 3.24 | 0.001346 |
| Timing | Ramping Pre-Tone | Intercept | 152 | 12.77 | 1.79 | 7.15 | 9.38E-11 |
| | | Log ($T_{tone}$) | | 0.67 | 0.20 | 3.41 | 0.000912 |
| EEG | N2 | Intercept | 85 | -0.12 | 0.02 | -5.47 | 4.50E-07 |
| | | P(Error \| $SS_{seen}$) | | 0.04 | 0.02 | 2.42 | 0.0178 |
| EEG | P3 | Intercept | 85 | 0.23 | 0.02 | 10.60 | 3.16E-17 |
| | | Log ($T_{tone}$) | | 0.04 | 0.01 | 3.72 | 0.000352 |

**Supplementary Table 2 | Comparison of model fits to different signals and periods**

**Top**, Heatmap of difference in BIC values (ΔBIC) for each model applied to the indicated neural signals and response periods. We fitted spiking activity or ERP polarization to models derived from each of the behavioral / task measures. We considered models based on response conflict, associated error likelihood, plus SSD and feedback tone time and their associated hazard rates. These behavioral and task measures could be correlated, however, random variations of each allowed for their differentiation. A set of mixed-effects models were constructed to assess the relationship between neural activity against each of the testing behavioral / task measures, and compared using Bayesian Information Criteria (BIC).

The model with the smallest BIC was endorsed as the best (black fill), and the relative performance of the other models is indicated by threshold ΔBIC values. Lower ΔBIC (red) indicates weaker support against other models, while higher ΔBIC (yellow) indicates stronger support against other models. Models that were not considered for a particular signal type or response period are denoted by crosshatch.

For Conflict neurons, the magnitude of post-SSRT modulation was best explained by the model with P(Error) as a predictor. This model obtained the lowest BIC compared to SSD or any other quantity, with weak support against the P(Error | $SS_{seen}$) (ΔBIC = 1.29) and strong support against other models (Δ BIC > 2.7). For Event Timing neurons, two time-periods were considered, because these neurons seem to modulate around two events during the task. Variation in the ramping activity before SSRT was best described by the time-based models of SSD with strong support against other models (Δ BIC > 2.7). The variation in dynamics of the ramping before the tone was best accounted for by the time of the feedback tone from the appearance of the stop-signal with strong support against other models (Δ BIC > 5.0). For Goal Maintenance neurons, the variation in magnitude of the phasic modulation was best accounted for by the log-transformed time of the feedback tone on canceled trials, with strong evidence against non-time-based models (Δ BIC > 3.0) and weak evidence against other time-based models (Δ BIC < 1). The variation in N2 amplitude was best accounted for by the P(Error | $SS_{seen}$) model (Δ BIC > 3.0 against competing models), with largest negativity when the experienced SSD was the earliest and associated with the lowest error likelihood. The variation in P3 amplitude was best accounted for by the log-transformed time of the feedback tone on canceled trials (Δ BIC > 4.0 against competing models) with weak support against other time-based models (Δ BIC < 1.30).

**Bottom**, Outcomes of mixed-effects modelling. Statistics are displayed for the model which best fits the neural modulation type and behavioral/task parameters. The mixed-effects model allowed for modeling random intercepts grouped by neuron (for spiking) or session (for ERP).

| Signal type | N | Proportion (%) | Conflict N | Conflict Proportion (%) | Event Timing N | Event Timing Proportion (%) | Goal Maintenance N | Goal Maintenance Proportion (%) |
|---|---|---|---|---|---|---|---|---|
| Error signal | 61 | 10.6 | 10 (5) | 13.3 | 25 (12) | 29.8 | 4 (1) | 7.4 |
| Gain signal | 91 | 15.8 | 11 (1) | 14.7 | 24 (9) | 28.6 | 4 (0) | 7.4 |
| Loss signal | 189 | 32.9 | 30 (4) | 40.0 | 14 (3) | 16.7 | 33 (1) | 61.1 |
| Other | 234 | 40.7 | 32 | 42.7 | 33 | 39.3 | 15 | 27.8 |
| Total Count | 575 | | 75 | | 84 | | 54 | |
| X2(3, N = 575) | test statistics | | 1.02 | | 44.86 | | 19.43 | |
| | p-value | | 0.79 | | **< 10-5** | | **0.00022** | |

**Supplementary Table 3 | Multiplexed signals between Error, Gain, and Loss with Conflict, Event Timing, and Goal Maintenance.**

Count of Conflict, Event Timing, and Goal Maintenance signals across the 575 recorded neurons relative to previously described Error, Gain and Loss[16]. The bottom row shows the test statistics for homogeneity based on a chi-square test. For each signal reported here, a 4-by-2 contingency matrix was constructed based on the counts of previously described neurons conveying the given signal. Conflict neurons multiplexed error, gain, and loss signals in proportion to their incidence of sampling. In contrast, Event timing neurons were significantly less likely to signal loss and more likely to signal error and gain than predicted by chance. Also, Goal Maintenance neurons were significantly more likely to multiplex with the loss signal than predicted by chance.
